# Sox2 and Sox3 are essential for development and regeneration of the zebrafish lateral line

**DOI:** 10.1101/856088

**Authors:** Cristian A. Undurraga, Yunzi Gou, Pablo C. Sandoval, Viviana A. Nuñez, Miguel L. Allende, Bruce B. Riley, Pedro P. Hernández, Andres F. Sarrazin

## Abstract

The recovery of injured or lost sensory neurons after trauma, disease or aging is a major scientific challenge. Upon hearing loss or balance disorder, regeneration of mechanosensory hair cells has been observed in fish, some amphibians and under special circumstances in birds, but is absent in adult mammals. In aquatic vertebrates, hair cells are not only present in the inner ear but also in neuromasts of the lateral line system. The zebrafish lateral line neuromast has an almost unlimited capacity to regenerate hair cells. This remarkable ability is possible due to the presence of neural stem/progenitor cells within neuromasts. In order to further characterize these stem cells, we use the expression of the neural progenitor markers Sox2 and Sox3, transgenic reporter lines, and morphological and topological analysis of the different cell types within the neuromast. We reveal new sub-populations of supporting cells, the sustentacular supporting cells and the neuromast stem cells. In addition, using loss-of-function and mutants of *sox2* and *sox3*, we find that the combined activity of both genes is essential for lateral line development and regeneration. The capability of sox2/sox3 expressing stem cells to produce new hair cells, hair cell-precursors, and supporting cells after damage was analyzed in detail by time-lapse microscopy and immunofluorescence. We are able to provide evidence that sox2/3 expressing cells are the main contributors to the regenerated neuromast, and that their daughter cells are able to differentiate into most cell types of the neuromast.

## INTRODUCTION

Sensory systems are susceptible to damage due to exposure to the external environment. As in other tissues, sensory systems vary in their capacity for cell renewal and self-repair among vertebrate species (Bermingham-McDonogh and Reh, 2011). For instance, mammals have a limited capacity for regenerating many tissues. In contrast, non-mammalian vertebrates show an extraordinary regenerative capacity in most tissues and organs (Borchardt and Braun, 2007; Kaslin et al., 2008; Brignull et al., 2009; reviewed in Bonfanti, 2011).

In mammals, vestibular hair cells of the inner ear have a limited capacity for regeneration (Forge et al., 1993; Kawamoto et al., 2009; Burns et al., 2012; Golub et al, 2012), and auditory hair cells cannot be replaced after birth (White et al., 2006; Cox et al., 2014; Burns and Corwin, 2013). In contrast, in birds, fish and amphibians, hair cells regenerate constantly, during their entire life cycle (Cotanche, 1987; Balak et al., 1990; Baird et al., 1993; Edge and Chen, 2008; Rubel et al., 2013). In birds, it has been demonstrated that regeneration of new hair cells occurs via two mechanisms: by cell division of supporting cells (Corwin and Cotanche, 1988; Ryals and Rubel, 1988) or by transdifferentiation of supporting cells (Adler and Raphael, 1996; Robertson et al., 1996, 2004). In zebrafish, studies of the inner ear have shown that hair cell regeneration involves transdifferentiation of supporting cells in the absence of mitotic division in a *sox2*-dependent manner (Millimaki et al., 2010). Sox2 is a member of the SoxB1 subgroup of the Sox transcription factor family. SoxB1 proteins (Sox1, Sox2 and Sox3 in mammals) are neural-specific transcription factors largely known to maintain stemness of neural progenitors during vertebrate nervous system development (Graham et al., 2003; Sarkar and Hochedlinger, 2013). These proteins are involved in cell fate decisions during early development, as well as in neurogenesis, cell division and maintenance of adult neural stem cells (Mizuseki et al., 1998; Zygar et al., 1998; Buescher et al., 2002; Bylund et al., 2003; Kan et al., 2004; Miyagi et al., 2004). Sox2 is also required for the development of the mammalian inner ear. In this tissue, lack of Sox2 leads to failure of sensory epithelium specification and of hair and supporting cell generation (Kiernan et al., 2005). Hair cells are specified in all vertebrates by the transcription factor Atonal homolog 1 (Atoh1); a protein that counteracts Sox2 by inhibiting proliferation and promoting differentiation (reviewed by Atkinson et al., 2015).

Previous work has shown that Sox2 is expressed in the zebrafish lateral line (Hernández et al., 2007, Germana et al., 2009, Jiang et al., 2014), an exceptional model to study hair cell regeneration due to its simplicity and easy accessibility (Lush and Piotrowski, 2014). This sensory system is distributed throughout the surface of fish and allows for the detection of local water movements around their body (Ghysen and Dambly-Chaudière, 2007). The posterior lateral line is initiated by the migration of the posterior lateral line primordium from near the ear towards the tip of the tail. During its migration, cell clusters within the primordium start to differentiate, forming the protoneuromast. Protoneuromasts are deposited periodically and further differentiate into neuromasts, the individual organs of the lateral line. It has been described that Wnt and FGF signaling coordinate cell fate and morphogenesis in the migratory primordium, but the role of SoxB1 genes in these processes remains unknown. Each neuromast is comprised of mechanosensory hair cells and accessory cells of at least two known types: supporting cells, that underlie and surround hair cells; and mantle cells, located at the periphery forming a conical structure that covers the neuromast (Metcalfe et al., 1985; Ghysen and Dambly-Chaudière, 2007). The cell type diversity and architecture of the neuromast is still unclear, as some reports using novel genetic tools and fate mapping suggest that this organ comprises more than one supporting cell type (Romero-Carvajal et al., 2015; Viader-Llargués et al. 2018; Lush et al. 2019; Thomas and Raible, 2019). The lateral line of zebrafish exhibits a strong ability to regenerate mechanosensory hair cells after exposure to ototoxic compounds or heavy metals (Williams and Holder, 2000; Harris et al., 2003; Hernández et al., 2006; Ma et al., 2008), however, mechanisms and cell types involved in these processes remain still poorly identified. We have previously described the expression of Sox2 in neuromast supporting cells, which proliferate during regeneration, suggesting that these cells are the source of new hair cells (Hernández et al., 2007). However, the role of Sox2 and other SoxB1 members in lateral line development and regeneration remains unknown.

Here, using transgenic reporter lines combined with confocal imaging and 3D reconstruction, we establish an architectural map of the neuromast in which sustentacular supporting, and stem cells, represent two functionally and spatially different cell types within the neuromast. The pool of progenitor cells that express Sox2 and Sox3 undergoes cell renewal and gives rise to new cell types within regenerated neuromasts. In contrast, sustentacular supporting cells neither proliferate nor originate new cell types. In addition, these cells surround hair cells basally and laterally, suggesting a structural and functional role in the neuromast. By knock-down and loss-of function approaches, we show that *sox2* and *sox3* are important for the development of the lateral line, both for the protoneuromast formation and timing of pro-neuromast deposition during migration of the primordium, and for the number of hair cells in the neuromasts. Furthermore, we reveal that *sox2* and *sox3* allowing the cells proliferation in the neuromast and would be required for hair cell regeneration. Our work provides novel insights into the topological organization of highly regenerative peripheral sensory organs, revealing details of the relationship of regenerating neurons with their niche.

## MATERIALS AND METHODS

### Zebrafish Maintenance

Zebrafish were maintained at 28.5°C on a 14 h light/10 h dark cycle. Fish were kept under standard conditions as described by Westerfield, (Westerfield, 2007). Embryos were raised in E3 media (Westerfield, 2007) in a 28.5°C incubator. The following zebrafish strains were used: AB strain wild-types (Zebrafish International Resource Center, Eugene, OR); *Tg(pou4f3:gap43-GFP)_s356t_*, which expresses GFP under the control of the Brn3c promoter, in the hair cell membrane (Xiao et al., 2005); *Tg(Bglobin/SOX3_HEs8:GFP)*, which expresses GFP in the Sustentacular Support Cells (SuSC) as we describe here, made from a 1kb highly conserved noncoding element (HCNE) upstream of the human SOX3 gene (Navratilova et al., 2009; obtained from the lab of Dr. Thomas Becker); *Tg(atoh1a:dTomato)*, which expresses RFP in the posterior lateral line (pLL) unipotent hair cell-precursors (Kani et al., 2010, obtained from the lab of Dr. Shin-ichi Higashijima); *Tg(-8.0cldnb:lynEGFP)_zf106_*, which express GFP in the membrane of all cells of the lateral line system (Hass and Gilmour, 2006, obtained from ZIRC); *Et(krt4:EGFP)sqet20*, which express GFP in the mantle cells of the lateral line (Parinov et al., 2004, obtained from ZIRC); and the *sox2* and *sox3* mutant lines *sox2_x50_*, *sox3_x52_* (Gou et al., 2018a). For live imaging, embryos were anesthetized with Tricaine (3-amino benzoic acid ethylester; Sigma), 4 mg/ml in E3 media and mounted in 0.5% low-melting agarose (Sigma) in E3 media.

### Copper sulfate treatment

To damage the different cell types present in the posterior lateral line of zebrafish, larvae were incubated in different CuSO_4_ concentrations (Hernández et al., 2006). CuSO_4_ (Merck, Darmstadt, Germany) stock was prepared in bi-distilled water (Winkler, Santiago, Chile), and the different dilutions were made in E3 media. Larvae were incubated in Petri dishes for the appropriate times at 28.5°C, rinsed three times with E3 media and embryos were fixed with 4% paraformaldehyde in phosphate buffered saline (PFA-PBS) overnight at 4°C. For the regeneration cycle experiments, the *Tg(pou4f3:gap43-GFP)_s356t_* larvae were incubated in 10 μM CuSO_4_ for 1 h, and the number of hair cells were counted every 24 or 48 h. The treatment was repeated for four cycles of injury and regeneration.

### BrdU Treatment

To label cells in S-phase, larvae were treated with pulses of 10 μM BrdU (Merck, Darmstadt, Germany, dissolved in E3) for various lengths of times depending on the experiment, as indicated in the text. Embryos were fixed in 4% PFA-PBS overnight at 4°C. BrdU immunostaining was performed as described below.

### Whole mount Immunocytochemistry and *in situ* hybridization

Larvae were fixed in 4% PFA overnight at 4°C and then rinsed in PBST (phosphate buffer saline with 0.1% Tween-20). Fish were blocked in 2% goat serum, 1% BSA, 1% DMSO and 0.1% Triton X100 for 1 h and later incubated with different antibodies overnight at 4°C (Primary antibodies: anti-Sox2 1:200, Chemicon; anti-Sox3 1:200, a kind gift from Dr. S. Wilson; anti-SoxB1 1:500, a kind gift from Dr. H. Kondoh; anti-BrdU 1:500, Dako; anti-acetylated Tubulin 1:1000, Sigma). Larvae were rinsed in PBST three times for 20 min and incubated overnight at 4°C with secondary antibodies (anti-mouse 1:500 IgG-Alexa Fluor 488, 1:500 IgG Alexa Fluor 594, Molecular Probes; anti-rabbit IgG-Alexa Fluor 488, 1:500 IgG-Alexa Fluor 594; 1:500 IgG-Alexa Fluor 633, Molecular Probes). The larvae were rinsed with PBST three times for 20 min and mounted for imaging. For labeling of cells that had incorporated BrdU, fixed fish were treated with proteinase K for 20 min, re-fixed in 4% PFA, then treated with 2N hydrochloric acid for 1 h, rinsed with PBST, blocked and continued with the normal immunocytochemistry protocol mentioned above.

*In situ* hybridization was done following the protocol of Thiesse et al., 2008. Digoxygenin-labeled antisense RNA probes against *deltaB*, *notch3* and *atoh1a* were hybridized to fish at 28, 34 and 36 hpf. Detection was via alkaline phosphatase-conjugated anti-digoxygenin (Sigma), followed by NBT/BCIP color reaction (Boehringer Mannheim) and stopped by three rinses in PBST.

### DAPI staining

Embryos were fixed in 4% PFA at 4°C overnight and then washed in PBST three times, PBS two times before incubating in PBS with 1µg/ml DAPI (Sigma D9542, stock prepared in DMSO) solution for 10 minutes in darkness. Stained embryos were washed at least three times in PBS before imaging.

### Morpholino knockdowns

Morpholinos and rhodamine-tagged Morpholinos were obtained from Gene Tolls (Corvallis, Oregon). The following morpholino sequences were used: *sox2,* 5’-AACCGATTTTCTCGAAAGTCTACCC-3’; *sox3,* 5’-CGTTTTCTTTCGAGTGCTTGGCACC-3’; and *sox2*/*sox3,* 5’-GCTCGGTTTCCATCATGTTATACAT-3’. Control morphants correspond to sibling embryos injected with equal doses of a scrambled control morpholino, and for the electroporated larvae, *cxcr4* MO (5’-ATGATGCTATCGTAAAATTCCATTT-3’) was used as a control.

### Larvae electroporation

Morpholino electroporation was performed in the transgenic lines *Et(krt4:EGFP)sqet20* and *Tg(-8.0cldnb:lynEGFP)_zf106_* at 72 hpf. The larvae were mounted in 0.05% agar dissolved in E3 and electroporated in the neuromast with M’s for *sox2*, *sox3*, *sox2/sox3* and *cxcr4* as a control. All morpholinos were labeled with rhodamine for electroporations and to confirm the incorporation of the M’s in the neuromast cells. After the procedure, the larvae were left in E3 medium for at least 1 h, and later incubated with copper solution to eliminate the hair cells. Larvae were analyzed 30 h after copper incubation to count the number of rhodamine labeled cells.

### Image acquisition and time-lapse imaging

All images were taken in a Zeiss LSM 510 or LSM 710 confocal microscope, a Zeiss Axiovert 200M fluorescence microscope, or an Olympus IX81 fluorescence microscope, using 20x air, 40x air and 65x oil objectives. All images were processed using ImageJ software. For time-lapse imaging, the fish were anaesthetized in 0.01% tricaine (Sigma) and mounted in 1.5% low melting point agarose (Winkler) dissolved in E3. Movies were recorded in an Olympus IX81 fluorescence microscope and processed using ImageJ software. Images were acquired every 40 min for 8 or 12 h.

### Statistical analysis

All experimental data are expressed as mean +/- SEM. The data obtained were analyzed using GraphPad Prism software (GraphPad Software, La Jolla, CA, USA) and compared using unpaired two-sided Student’s t-test and one-way ANOVA. The probability level for statistical significance was P < 0.05.

## RESULTS

### Distribution and proliferation of cells expressing Sox2 and Sox3 in the zebrafish Posterior Lateral Line

We first aimed to identify cells expressing Sox2 and Sox3 within the posterior lateral line (pLL) system. Using specific antibodies for both proteins, we performed immunofluorescence at different stages of LL development. In line with previous reports (Hernández et al., 2007; Jiang et al., 2014; Steiner et al., 2014), we observed Sox2 expression in the majority of cells within the pLL migratory primordium, in neuromasts and interneuromastic cells; a similar pattern was found for Sox3 (**Figure S1A-E**). We then focused on identifying cells expressing Sox2 and Sox3 within neuromasts, including sensory hair cells and accessory cells (defined as all non-sensory hair cells within neuromast). These proteins were rarely detected in mature hair cells (<7% of co-localization, data not shown), as evidenced in *Tg(brn3c:mGFP)* transgenic larvae (**Figure S1D-D’’**). Accessory cells include supporting cells underlying and laterally surrounding hair cells, and mantle cells located at the apical periphery of neuromasts, forming a conical structure that covers the neuromast (Ghysen and Dambly-Chaudière, 2007). By using the transgenic line *Tg(HEs8:GFP)*, we found that supporting cells are composed of at least two populations with different morphologies and location with respect to hair cells and mantle cells (Navratilova et al., 2009). Fish harboring this transgene express GFP under the control of a human SOX3 enhancer and we used GFP expression, combined with morphological recognition and immunostaining against Sox2 or Sox3, to distinguish between the two cell types; we note that we do not consider GFP expression to be a reporter of the endogenous expression of zebrafish *sox3* (**Figure 1A-F**). We found high GFP expressing cells (*Hes8:GFP_hi_*) in close contact and completely surrounding hair cells (**Figure 1C-D’’** and **Movie S1** and **S2**) while only some of these cells expressed Sox2 and Sox3. In contrast, low GFP expressing cells (*Hes8:GFP_low_*) located basally and peripheral to hair cells consistently expressed Sox2 and Sox3 (**Figure 1G-K’’** and **Figure S1F-H’’** for Sox2, and **S1H-I’’** for Sox3). We noticed that, within the basal cell layer of the neuromast, Sox2 and Sox3 expression was detected in rounded cells located at the center and at the periphery, whereas at the apical level they were localized only along the contour of the neuromast and not in the center, where hair cells reside (**Figure S1B-B’’** and **S1F-G’’** for Sox2, **Figures S1D-E’’** and **S1I-J’’** for Sox3) further suggesting that this could be a new population of accessory cells. It has been previously proposed to name all supporting cells as Sustentacular Supporting Cells (SuSCs) (Pinto-Texeira et al., 2015). However, we observed that supporting cells are a heterogenous population, comprising cells with different morphology and localization with respect to hair cells. Therefore, we made use of the name “Sustentacular” to refer only the supporting cells showing a central location, a drop-like shape, and completely surrounding and holding hair cells.

**Figure 1.**
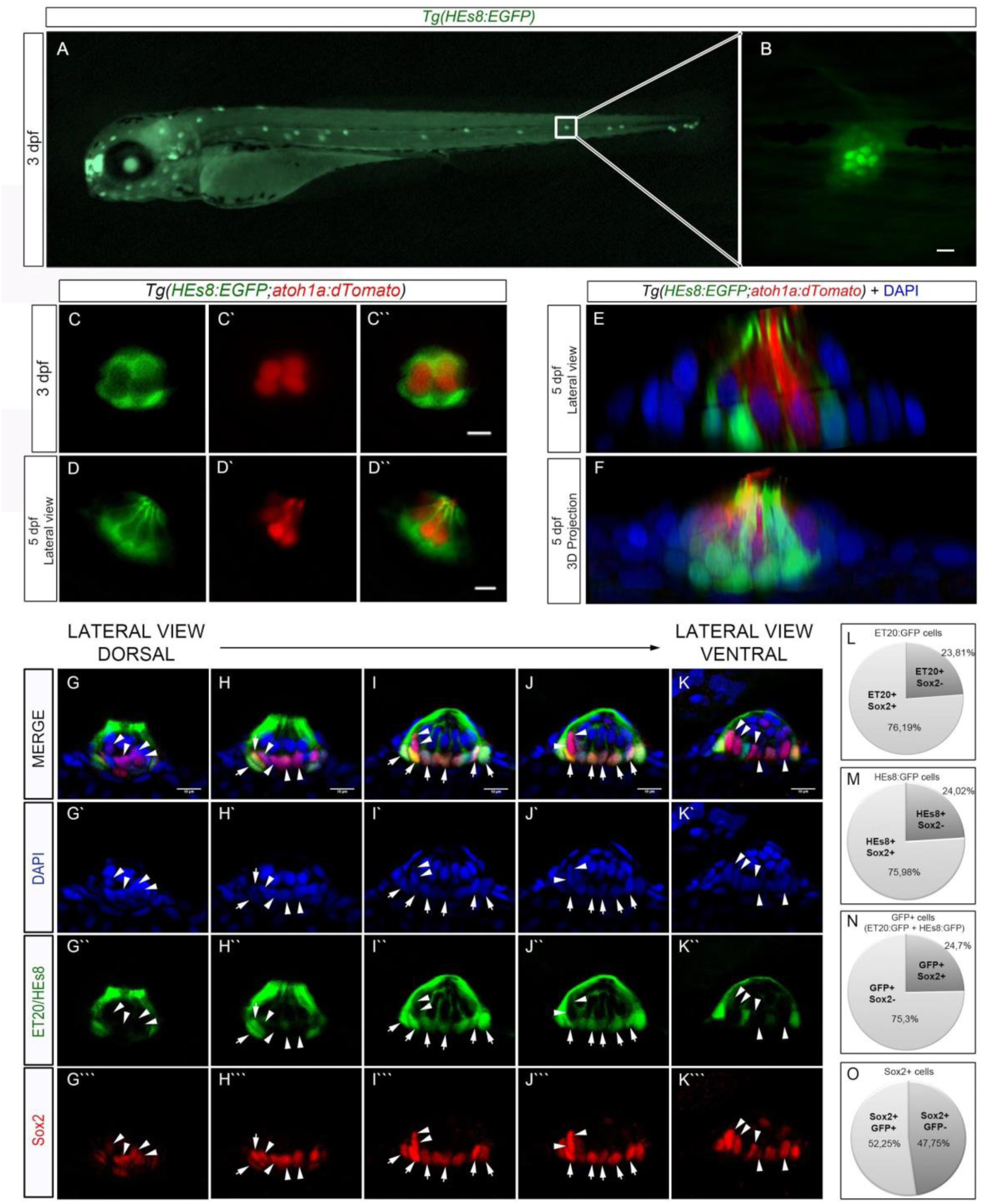
Specific neuromast cell type markers. (A,B) *Tg(HEs7:EGFP)* transgenic line expressing green fluorescence in lateral line neuromasts. (C-F) Confocal images of a double transgenic line *Tg(HEs7:EGFP;atoh1a:dTomato)* showing Sustentacular Support Cells (SuSCs, labeled in green, C,D) and Unipotent Hair Cell-Precursors (UHCP, labeled in red, C’,D’) at 3 dpf. Note the absence of colocalization when merged (C’’,D’’). (E,F) Neuromast lateral views of a double transgenic line *Tg(HEs7:EGFP;atoh1a:dTomato)* at 5 dpf. DAPI was added to visualize the nuclei. (E) Lateral reslices of a serial stack image. (F) 3D reconstruction of another serial stack image. (G-K’’’) Neuromast lateral views of 4 dpf double transgenic larvae *Tg(ET20:EGFP;HEs7:EGFP)* immunostained for Sox2. Confocal serial stacks from dorsal (left) to ventral (right). Arrowheads: Sox2 staining without GFP colocalization. Arrows: Sox2/GFP colocalization. (L-O) Sox2 labelled cell number quantification with respect to the other cell markers: Mantle cells (L), Sustentacular support cells (M) and both SuSC and Mantle cells (N,O). Note that in (N) Sox2 positive cells are measured with respect to the total number of SuSC and Mantle cells but in (O), the SuSC and Mantle cells are quantified with respect to the total number of Sox2 labelled cells. Scale bars = 10 μm.

The Sox2_+_ and Sox3_+_ cells that are not SuSCs could be mantle cells. Therefore, to distinguish them from known accessory cell populations, we performed Sox2 immunofluorescence in the *Tg(ET20:GFP)* line (Parinov et al., 2004) to visualize mantle cells, and in the *Tg(HEs8:GFP)* line to visualize SuSCs, and we quantified colocalization (**Figure 1G-K’’’**). We found that most mantle cells co-localized with Sox2_+_ cells (76 ± 5 %; n=20), similar to SuSCs (76 ± 6 %; n=7). That is, approximately 75 % of accessory cells co-expressed Sox2, both when they were counted separately (**Figure 1L,M**), and when they were counted together in the double transgenic *Tg(HEs8:GFP/ET20:GFP)* (**Figure 1N**). However, 52 ± 3.9 % of cells expressing Sox2 also expressed GFP, and almost half of the Sox2_+_ cells (48 %) do not correspond to any of the known accessory cell types (**Figure 1O**). We thought that these Sox2_+_, rounded cells located preferentially at the base of the neuromast, that are morphologically different from SuSCs and mantle cells (**Fig S2**), could be Neuromast Stem Cells (NmSCs, arrowheads in **Figure 1G-K’’’** and **S2A-E’’’**) and are potentially responsible for the regeneration of both, accessory and sensory hair cells (see below).

In this regard, we wondered whether this cell population is mitotically active from the onset of Sox2/Sox3 expression during primordium migration and in maturing neuromasts. When we exposed 30 hpf embryos to a 2 h pulse of BrdU, a substantial number of cells within the primordium had BrdU label co-localizing with Sox2 and Sox3 immunostaining (**Figure S3C-C’’, E-E’’**). On the other hand, only few cells within neuromasts were labeled with BrdU after applying an identical 2 h BrdU pulse in 3 dpf control larvae, although all of them expressed Sox2 or Sox3 (**Figure S3D-D’’, F-F’’**). Since proliferative cells are also positive for Sox2/Sox3, we evaluated the self-renewal capacity of these cells in the neuromast. We exposed 3 dpf larvae to 2 h pulses of BrdU and we evaluated BrdU labeling plus phospho-histone H3 immunostaining (mitosis specific marker; PH3) after 24, 48 and 72 h post pulse (chase, **Figure S3G**). We found, in all chase periods analyzed, that most BrdU_+_ cells were also PH3_+_ (**Figure S3H**, red bars), indicating that the initial mitotic activity of Sox2/Sox3_+_ cells is maintained over time, suggesting that these cells could be self-renewing. Even when we analyzed the neuromasts of 3 dpf larvae exposed to 24 h BrdU pulses one week later, we found that more than 40% of their cells were double labeled with both markers, BrdU and PH3 (**Figure S3H**, blue bars). This result supports the notion that Sox2/Sox3_+_ cells self-renew, an important characteristic of stem cells.

Next, we evaluated whether cells that incorporated BrdU in the migratory primordium are later found within neuromasts. We determined the number and final position of Sox3 positive cells that incorporated BrdU during primordium migration (30 hpf; 2 h BrdU pulse) after 48, 72 and 96 h post-pulse (hpp; **Figure S4A**). Most BrdU_+_ cells were found in the periphery of the neuromast at 48 hpp (**Figure S4C-C’’**), which later shifted towards a more central position (**Figure S4D-E’’**). Moreover, the percentage of BrdU_+_ cells that co-stained with anti-Sox3 increased from ~75% at 48 hpp to 95% at 96 hpp (**Figure S4B**), indicating that proliferative cells in the migrating primordium give rise to nearly all Sox3_+_ cells within mature neuromasts. To evaluate if the Sox3_+_ cells could proliferate and maintain this capacity after damage, we analyzed the behavior of the proliferative population in steady state and after a copper damage. We performed a 2 h BrdU pulse followed by 1 h of 10 μM copper sulphate treatment (to eliminate only hair cells) and evaluated BrdU incorporation and Sox3 expression at 24, 48 and 72 h post-pulse (**Figure S4F**). We observed a constant percentage of Sox3_+_ cells (roughly 30%) that proliferated and maintained the BrdU label after a damage-regeneration process at each time-point (**Figure S4G**). These data show that there is a population of progenitors in neuromasts, which in normal conditions proliferate, and that after damage, they self-renew while contributing to the regeneration process. In this regard, Sox3-positive cells behave similarly to the Sox2-positive cells and could label the same population of neuromast stem cells (NmSCs).

### Both SuSCs and unipotent hair cell-precursor cells originate from multipotent Sox2+ progenitor cells

Previous work has shown that the *atoh1a* gene is expressed in unipotent hair-cell progenitors, UHCPs (Wibowo et al., 2011), just before they give rise to two hair cells by symmetric cell division (Video S3). Since Sox2_+_ cells self-renew and new hair cells regenerate from *atoh1a* UHCP cells, we wondered whether other cell types within neuromasts (*i.e.*, SuSCs and UHCPs) could also regenerate from this pool of progenitors. To address this, we incubated *Tg(HEs8:GFP)* and *Tg(atoh1a:tdTomato)* larvae with 10 μM CuSO_4_ for 2 h in order to ablate all hair cells, UHCPs and SuSCs (Hernández et al., 2007). Subsequently, we fixed larvae 24 and 48 h after CuSO_4_ treatment and we evaluated Sox2 protein co-expression with each transgenic marker (**Figure 2A**). While around 75% of *HEs8* and *atoh1a+* cells co-localize with Sox2 under control conditions (**Figure 1M** and data not shown), after damage co-localization of these markers increases to 100% (**Figure 2B**). This result suggests that both new SuSC (*GFP*_+_) and new UHCPs (*tdTomato*_+_) were regenerated from Sox2_+_ progenitor cells (**Figure 2C-D’’** and **2E-F’’**, respectively). These results reinforce the hypothesis that Sox2-positive cells give rise to different neuromast cell types after injury.

**Figure 2.**
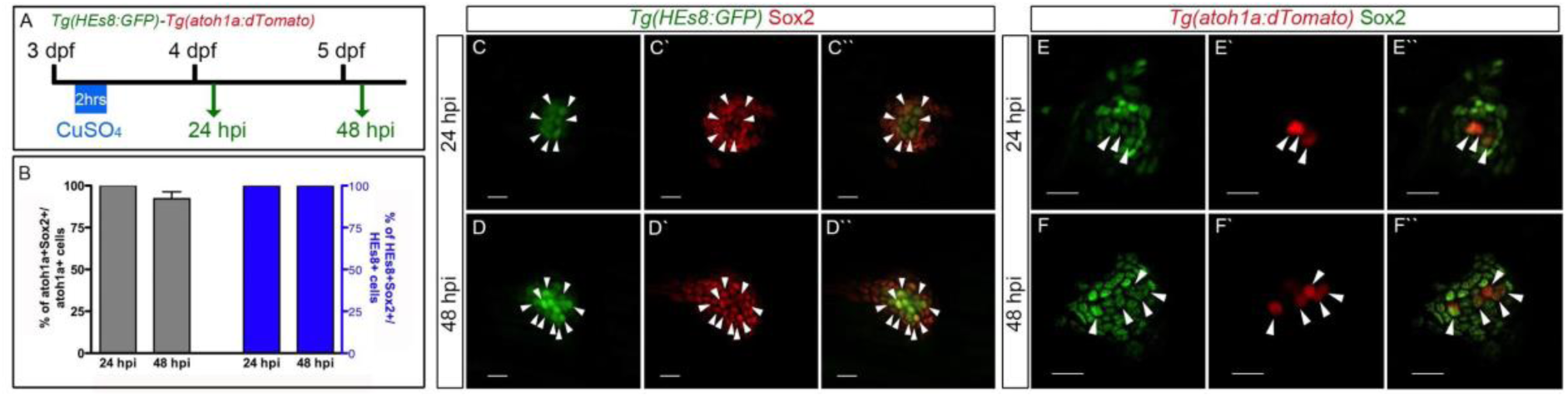
New sustentacular support and hair cell-precursor cells come from Sox2-positive progenitors. (A) *Tg(HEs8:GFP)* or *Tg(atoh1a:dTomato)* 3 dpf larvae were incubated in 10 μM copper sulphate for 2 h (eliminating both hair cells and sustentacular support cells) and fixed for Sox2 antibody staining at 24 and 48 hour post incubation (hpi). (B) Sox2 labelled cells measurement with respect to the total number of *HEs8:GFP* (left) and *atoh1a:dTomato* (right) marked cells after CuSO_4_ treatment. (C-F’’) Confocal images showing Sox2 colocalization (arrowheads) with *HEs8:GFP* (C-D’’) or *atoh1a:Tomato* (E-F’’) after 24 (C-C’’; E-E’’) and 48 (D-D’’; F-F’’) hours after damage. Scale bars = 10 μm.

Since Sox2 colocalization with *atoh1a:tdTomato*_+_ cells was not seen in undamaged neuromasts (data not shown), or did not completely colocalize with all the SuSCs (*HEs8*_+_ cells, **Figure S1F-G’’**), we hypothesized that regenerated cells, shown in **Figure 2**, correspond to cells that had recently differentiated from a Sox2_+_ progenitor. Further, we found that 12 h and 24 h after CuSO_4_-induced damage, 100% of the remaining *HEs8:GFP* cells that survived the damage were also positive for Sox2 (**Figure 2B**). Therefore, we hypothesized that the remaining *HEs8:GFP*_low_ cells left after damage were *HEs8*_+_ cells that normally reside in basal and peripheral positions in the neuromast, express Sox2 or Sox3 (**Figure S1F-J’’**), display a rounded shape, and express low levels of GFP. These cells are clearly distinct from the SuSCs, which exhibit high levels of GFP and that form a tight interaction with hair cells surrounding them. Since the *HEs8_+_/GFP*_low_ cells left after neuromast damage could recapitulate the behavior of Sox2_+_ progenitors in a regenerative context, we followed their dynamics in time-lapse imaging experiments. For this, we analyzed *Tg(HEs8:GFP/atoh1a:tdTomato)* double transgenic larvae after 2 h of 10 μM CuSO_4_ treatment (this protocol eliminates hair cells, UHCPs and SuSCs while maintaining the *HEs8_+_/GFP*_low_ cells, **Figure 3A**). We recorded different cell transitions during neuromast regeneration. In the series shown in **Figure 3B**, we detected the origin of two double-labeled cells from a symmetrical division of a progenitor cell expressing both markers. In another sequence (**Figure 3C**), after a symmetrical division of double-labeled cells, one of them lost GFP expression, divided again, and gave rise to two hair cell precursors (*HEs8:GFP*-/*atoh1a:tdTomato*_+_). Additionally, in one series (**Figure 3D**), we found two *HEs8:GFP*_+_ cells adopting the position of SuSCs, surrounding two *atoh1a:tdTomato*_+_ cells (left inset in **Figure 3D**). Altogether our data demonstrate the transition from multipotent Sox2_+_ progenitor cells to sustentacular supporting cells and unipotent hair-cell progenitors that differentiate into hair cells after mild damage.

**Figure 3.**
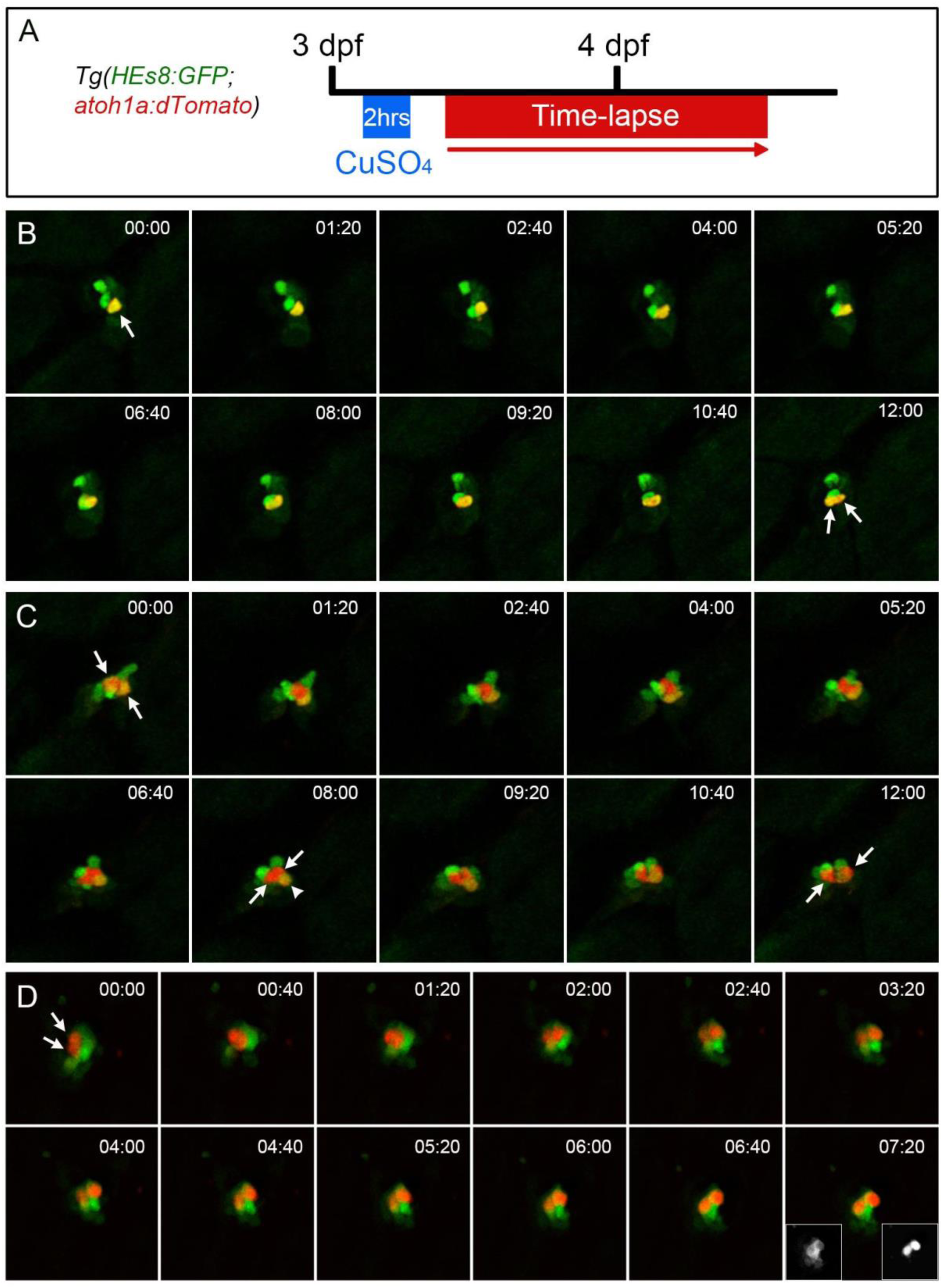
Transition from multipotent Sox2+ progenitor cells to unipotent hair cell progenitors and sustentacular support cells. (A) Double transgenic larvae *Tg(HEs7:GFP;atoh1a:dTomato)* were incubated 2 hours in 10 μM CuSO_4_ (eliminating hair cells, unipotent hair cell progenitors and sustentacular supporting cells) and time-lapsed after 1 hour post damage, where the first *atoh1a*+ cells appear. (B-D) Three different time-lapses showing the division of progenitor cells expressing both markers (B,C) and a hair cell progenitor (D). Pictures were taken every 40 minutes (B,C) or 80 minutes (D).

### Hair cells in the lateral line system continuously regenerate and induce overlapping of normally consecutive steps during forced regeneration

In order to further study Sox2_+_ cells and the regenerative capacity of neuromasts, we analyzed whether hair cell regeneration after copper damage is inexhaustible, as was shown previously in adult fish using Neomycin (Pinto-Texeira et al. 2015; Cruz et al. 2015). We exposed larvae repeatedly to CuSO_4_ to induce injury followed by removal of the metal to allow hair cell regeneration. When we incubated 3 dpf *Tg(brn3c:mGFP)* transgenic larvae with 10 μM of CuSO_4_ for 1 h, all hair cells in the pLL were eliminated (**Figure S5A,B**). As expected, during the next 24 h, new hair cells emerged (**Figure S5C**). When we exposed the same larvae to three successive rounds of exposure to 10 μM CuSO_4_ for 1 h each, all neuromasts regenerated their hair cells 24 h after each injury (**Figure S5D-I**). We quantified the number of regenerated *GFP*_+_ hair cells in the first five neuromasts of the pLL 48 h after each exposure, when a stable number of regenerated hair cells are reached (Hernández et al., 2006 and data not shown). The number of hair cells in 5 dpf non-treated larvae was similar to larvae exposed to one round of damage plus regeneration. In the following rounds of regeneration, the number of hair cells per neuromast decreased slightly after the second (73.2%, n=24) and third (71.2%, n=24) rounds of regeneration (in comparison to untreated larvae, see **Figure S5J**). This long-term hair cell regeneration assay demonstrates that lateral line neuromasts have a high capability of repeatedly regenerating hair cells after copper damage.

In order to test the proliferative potential of progenitor cells and the fate of the new cells produced in a context of repeated loss and regeneration of hair cells, we carried out pulse chase experiments with BrdU (**Figure 4A**). Again, we induced damage using a 1 h treatment with 10 μM of CuSO_4_ to eliminate hair cells followed by a 24 h BrdU-pulse (**Figure 4A**). We found that, immediately after BrdU pulse, about 60% of Sox2_+_ cells proliferate (**Figure 4B and F**). The proportion of Sox2_+_ cells that showed BrdU labeling increased when the larvae were then exposed to cycles of hair cell damage and regeneration every 24 h (**Figure 4C-F**). Further, 100% of BrdU_+_ cells expressed Sox2 at these time points (**Figure 4G**). This result indicates that Sox2_+_ cells are self-renewing and that they maintain Sox2 expression without losing the BrdU label, possibly due to a low rate of cell division. On the other hand, the Sox2_+_/BrdU_-_ cells that remained after the first CuSO_4_-induced damage but were not found in later cycles of damage/regeneration may have undergone differentiation (without proliferating), gradually diminishing in number (**Figure 4F**). To support this notion, we found that the proportion of regenerated hair cells (*brn3c*_+_) that came from BrdU_+_ progenitor cells increased after each CuSO_4_ treatment (**Figure 4H-L**). Strikingly, we also found that these new hair cells also expressed Sox2, which could be the result of the rapid succession of damage and regeneration cycles (**Figure 4M-P**). This condition is usually not observed in a control, undamaged, neuromast (Jiang et al., 2014) though it can be observed occasionally (Lush et al., 2019 and less than 7% from our data). When we quantified the number of *brn3c:GFP_+_*/Sox2_+_ cells in relation to the total number of *brn3c*_+_ hair cells, we found a progressive increase, reaching almost 100% of co-labeling at 72 h post BrdU pulse (hpp) and after three cycles of damage and regeneration (**Figure 4Q**). Possibly these highly demanding conditions for rapid hair cell regeneration generate a co-occurrence of both markers. These results show that, when the pLL system is forced to regenerate continuously, Sox2_+_ cells are able to differentiate and give rise to new hair cells, suggesting a high regenerative capacity driven by NmSCs.

**Figure 4.**
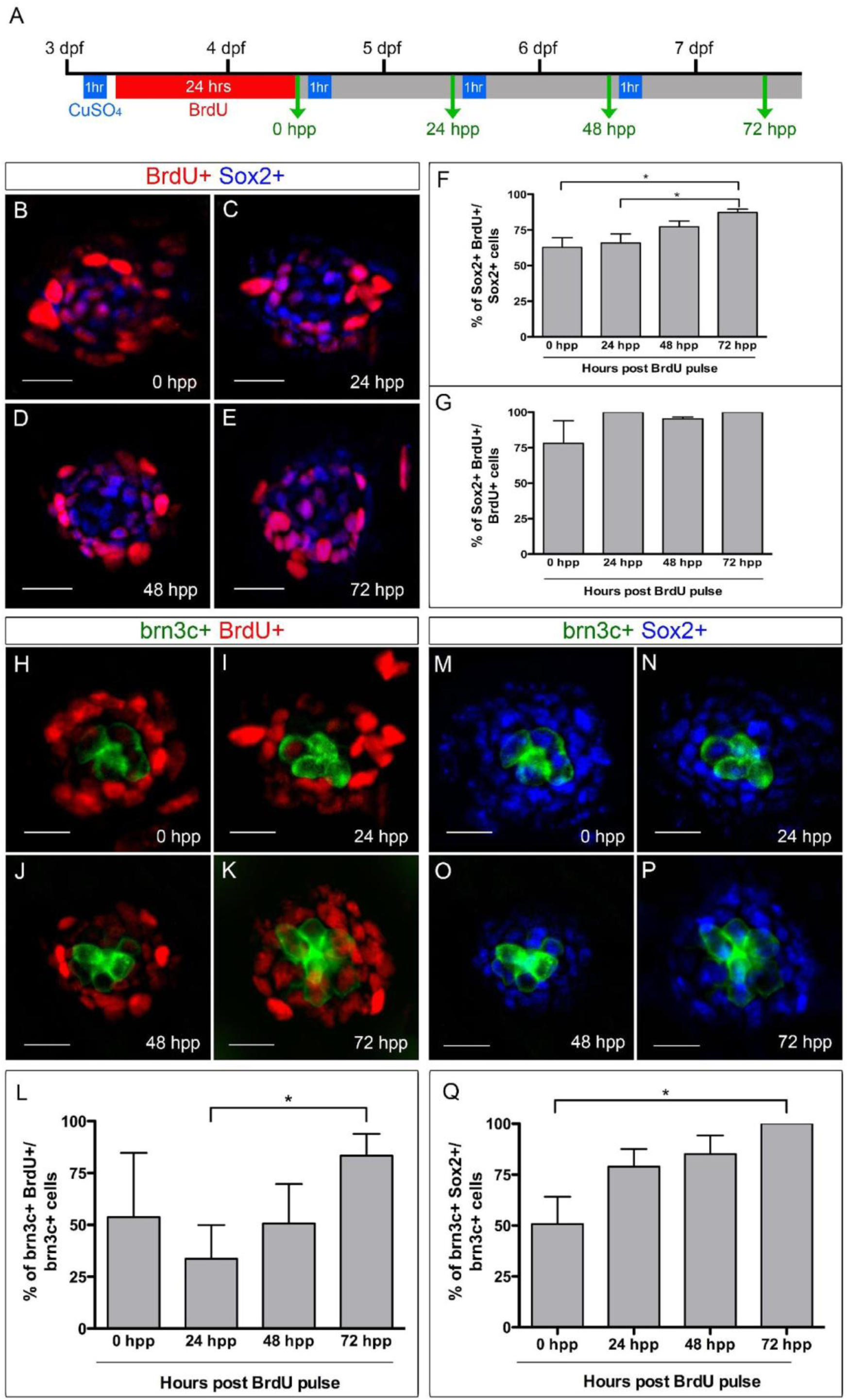
Sox2_+_ cells proliferate, self-renew and give rise to hair cells after successive rounds of regeneration. (A) Scheme representing the BrdU incubation for 24 hours performed after a first hair cell ablation at 3 dpf, followed by 3 successive hair cell ablations, one immediately after BrdU and the other two 24 and 48 hours later. This protocol forced neuromasts to be continuously regenerating causing an overlap between regeneration steps. The green arrows represent the time-points (hour post BrdU pulse, hpp) when treated larvae were fixed. (B–E) Confocal images of neuromasts showing Sox2_+_ cells (blue), BrdU_+_ cells (red) and double labeled cells (magenta) at different time-points after the BrdU pulse. (F-G) Quantification of double labeled cells relative to total number of Sox2_+_ cells (F) or total number of BrdU_+_ cells (G), at each time-point after BrdU pulse. (H–K) Confocal images of neuromasts showing regenerated hair cells (*Tg(brn3c:mGFP)* in green) and BrdU_+_ cells (red) at different time-points after the BrdU pulse. (L) Quantification of double labeled cells (hair cells coming from progenitors that proliferated during the BrdU pulse) relative to the total number of regenerated hair cells. (M–P) Confocal images of neuromasts showing regenerated hair cells (green) and Sox2_+_ cells (blue) at different time-points after the BrdU pulse. (Q) Quantification of double labeled cells (hair cells coming from progenitors that also express Sox2) relative to the total number of regenerated hair cells. P < 0.05. Scale bars = 10 μm.

### Loss-of-function of *sox2* and *sox3* genes impairs posterior lateral line formation and are necessary for proliferation in neuromasts

Both *sox2* and *sox3* are expressed in the pLL primordium and neuromasts (Hernández et al., 2007; Navratilova et al., 2009 and Figure S1). In order to evaluate whether *sox2* and/or *sox3* are involved in pLL development, we examined homozygous null mutants for both genes also carrying the *brn3c:mGFP* transgene (Gou et al. 2018a). Expression of the encoded reporter protein allowed us to count hair cells in each neuromast at 3 dpf. In addition, we used DAPI staining to label cell nuclei, facilitating the identification and position of neuromasts at 46 hpf. We found no statistically significant difference in the number and distribution of pLL neuromasts in *sox3* single mutant larvae compared with siblings at 46 hpf (**Figure 5 A,C**). However, we found a significant decrease in the average number of neuromasts between both *sox2* single mutants and *sox2/sox3* double mutants compared to control siblings; in both cases, the neuromasts were accumulated towards the posterior half of the trunk and tail (**Figure 5 B,D,F**). When we compared the number of hair cells in mutant and WT neuromasts, we found that *sox2* mutants have less hair cells compared to control siblings (P<0.01; **Figure 5 F,N**). Further, *sox2/sox3* double mutants showed a stronger reduction in the number of hair cells (P<0.0001; **Figure 5 F,N**). In addition, some of the *sox2* single and *sox2/sox3* double mutants also show a short body axis and an upward curled tail, as we have previously shown (Gou et al. 2018a). These data indicate that both *sox2* and *sox3* genes are important for protoneuromast formation and timing of pro-neuromast deposition during migration of the primordium, as well as to reach the correct number of hair cells within neuromasts.

**Figure 5.**
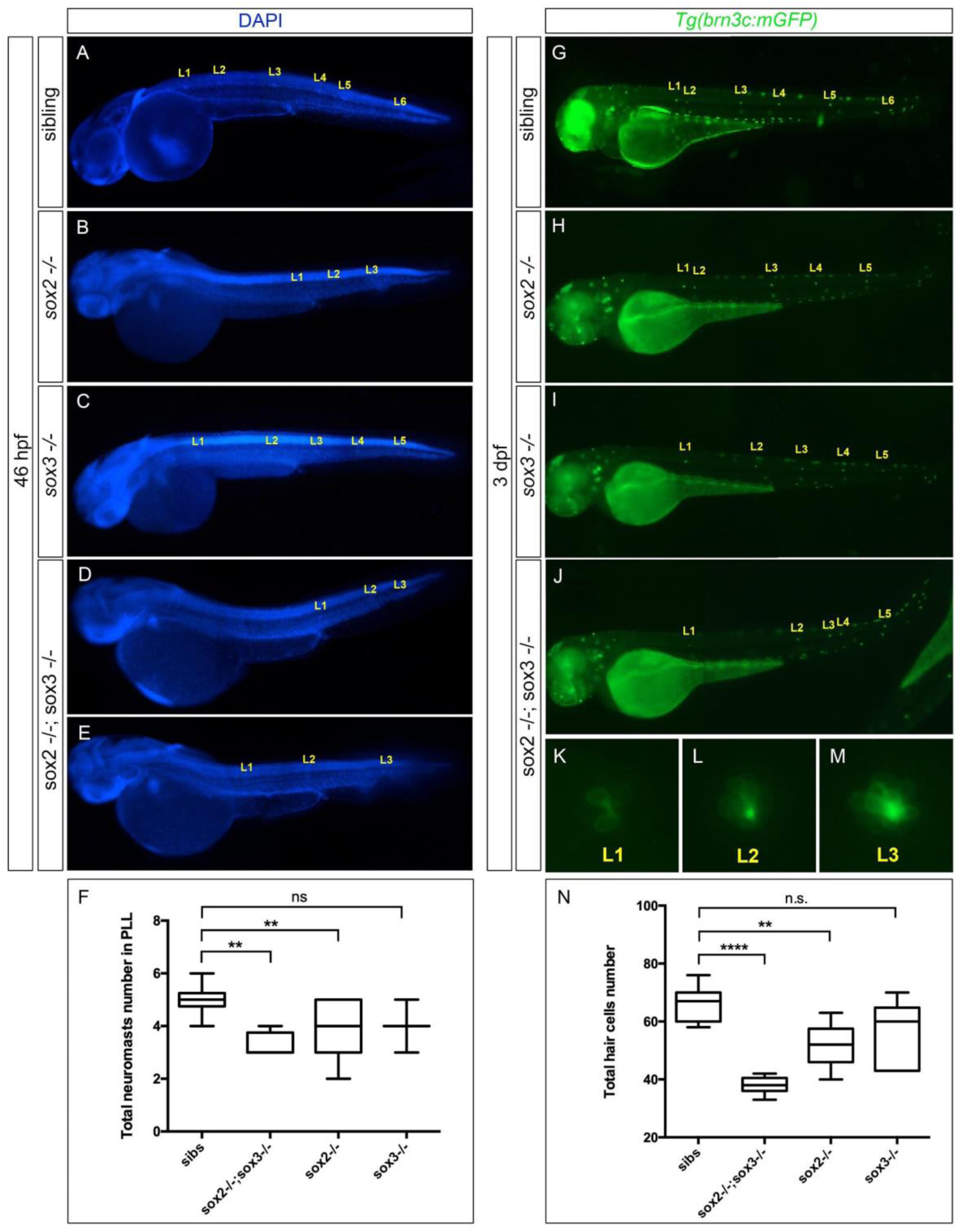
Major defects in the posterior lateral line of *sox2/sox3* double mutant larvae. (A) Sibling and (B,C) *sox2/sox3* double mutants larvae at 46 hpf labeled with DAPI to visualize deposited neuromasts. (D) Total number of pLL deposited-neuromasts in siblings (sibs), *sox2/sox3* double mutants and *sox2* and *sox3* single mutant larvae at 46 hpf. The *sox2* single mutants and the *sox2/sox3* double mutants differ from wt larvae in the number of deposited-neuromasts with p<0,01. (E) Sibling and (F-I) *sox2/sox3* double mutants transgenic larvae Tg(brn3c:mGFP) at 3 dpf. (G-I) Close up to neuromasts L1, L2 and L3 from a similar larva to the one in (F). (N) Total number of pLL hair cells in siblings (sibs), *sox2/sox3* double mutants and *sox2* and *sox3* single mutant larvae at 3 dpf. The *sox2* single mutant and the *sox2/sox3* double mutants differ from wt larvae with p<0,01 and p<0,0001 respectively. L1-L6: posterior lateral line neuromasts. n.s.: not significant; ** p<0,01; **** p<0,0001.

An additional tool to study gene function in the zebrafish embryo and larva is the use of gene knock-down by morpholino injection. Thus, to complement the mutant analysis, we injected 1-cell WT and *Tg(clndB:lynGFP)* embryos with antisense morpholino oligonucleotides designed to suppress the expression of *sox2*, *sox3* or both genes (*sox2*MO, *sox3*MO and *sox2/sox3*MO, respectively). We evaluated primordium migration, neuromast deposition and patterning, and hair cell differentiation in morphant fish. At 46 hpf, *sox2* and *sox3* morphant larvae showed no difference in neuromast number or position when compared to non-injected control embryos (**Figure S7A-B**). However, injection of the *sox2/sox3*MO affected neuromast positioning at 36 hpf (the most anterior pLL neuromasts were not present, see **Figure S6B**), though the total number of neuromasts remained unchanged. Thus, the correct number of proneuromasts were deposited by the migratory primordium, but deposition was delayed in mutants compared to control animals (**Figure S6C**). As was observed in the single and double mutants, *sox2*MO, *sox3*MO and *sox2/sox3*MO double morphants displayed a reduction in the total number of hair cells per neuromast at 3 dpf,; this difference was determined both by analyzing the phenotypes in the transgenic *Tg(brn3c:mGFP)* background, as well as by DiAsp staining of functional hair cells (Figure S7I-R). To complement our analysis of morphant animals, we carried out *in situ* hybridization using riboprobes complementary to mRNAs expressed in different cell types of neuromasts. Using probes against the hair cell markers *atoh1a* and *deltaB*, as well as *notch3*, which is expressed in supporting cell precursors (Itoh and Chitnis, 2001; Sarrazin et al., 2006; Matsuda and Chitnis, 2010), we confirmed the absence of neuromast deposition during the first half of primordium migration in *sox2/sox3* double knockdown fish (**Figure S6D-G** and **Figure S7C-H**). Additionally, double morphants showed no sign of protoneuromasts formation within the early migratory primordium (**Figure S6H-I**), had a significantly higher number of total cells in the migrating primordium at 28, 34 and 40 hpf (compared with the total cells in control larvae, see **Figure S6J**), as well as a significantly higher number of TUNEL positive cells at 34 hpf (but not at 28 hpf), compared to control larvae (**Figure S6K**). Therefore, our loss-of-function experiments show that *sox2* and *sox3* genes are essential for a proper development of the pLL in zebrafish.

While the analysis of *sox2* and *sox3* mutants and morphants demonstrates an important role for these genes in the development of the pLL neuromasts, it precludes the examination of their role in regeneration. To specifically address the role of *sox2* and *sox3* in hair cell regeneration, we decided to carry out loss-of-function experiments using neuromast electroporation of rhodamine-conjugated antisense morpholinos directed against these genes. For that, we used a morpholino specific to *cxcr4* (which is not expressed in the deposited neuromast; Ghysen and Dambly-Chaudière, 2007) as a control and we performed the electroporations in *Tg(clndB:lynGFP)* and *Tg(ET20:GFP)* 3 dpf transgenic larvae. Then, we electroporated control and *sox2/sox3* antisense morpholinos into fish that had been previously treated with a 2 h exposure to 10 μM CuSO_4_ in order to eliminate all hair cells and SuSCs (**Figure S8A**). When we compared the number of rhodamine_+_ cells (red) in neuromasts of larvae immediately after treatment (0 h post copper, hpc) and at 30 hpc, we found that fish electroporated with *cxcr4*MO (control) had an increase in labeled cells. Therefore, electroporated hair cell precursors or progenitor cells divide after hair cell elimination and contribute to neuromast regeneration (**Figure S8B,C,I,J,H,Q**). On the other hand, electroporation of *sox2*MO, *sox3*MO or *sox2/sox3*MO prevented these cells from undergoing cell division (**Figure S8D-H, K-Q**), a crucial step in the regeneration of hair cells from progenitor cells (Wibowo et al., 2011). Thus, in this context, our data shows that both *sox2* and *sox3* are required for the proliferation in the neuromast cells, an essential step to form new hair cells after damage.

## DISCUSSION

In this report, we show that cells present in the zebrafish lateral line neuromast are highly heterogeneous, in terms of molecular markers, shape, position, and behavior after injury. We also identify important functions for members of the SoxB family of transcription factors for development and regeneration of the zebrafish lateral line. Our work adds to the growing body of literature that has contributed to our understanding of the mechanisms of regeneration present in this mechanosensory organ, which can vary depending on the nature and magnitude of the injury. In the following paragraphs, we contextualize our findings with respect to previous work and we discuss the implications for further extending this research.

### Cell-specific markers distinguish different cell populations within lateral line neuromasts and their participation in the regenerative process

Studies performed in the avian cochlea have convincingly shown that supporting cells are the source of regenerated hair cells after their loss by chemical or physical injury (Corwin and Cotanche, 1988; Ryals and Rubel, 1988). Thus, characterization of supporting cells is essential to understand the mechanisms of non-mammalian hair cell regeneration. Here, we used the zebrafish *Tg(HEs8:GFP)* transgenic line to distinguish two types of supporting cell within neuromasts: i) differentiated Sustentacular Supporting Cells (SuSCs), which are central, beneath, and tightly associated with hair cells, or with committed unipotent hair cell-precursors, UHCPs, and ii) a population of cells that occupy a central and basal position within the neuromast, displaying a rounded shape that is distinct from that of SuSCs, and expressing the stem cell markers Sox2 and Sox3. Thus, the *HEs8* regulatory element, directs GFP expression to SuSCs, and to uncommitted progenitors, revealing the heterogeneity of the supporting cells in terms of their ability to differentiate into SuSCs, or hair cells in fish. In mammals, differentiated supporting cells are fate restricted in the cochlea and are completely unable to replace lost hair cells, though in the vestibular system they can contribute to a limited replacement of hair cells (Bucks et al., 2017). In the avian vestibular system, hair cells can be replaced -at a slow rate-by transdifferentiation of supporting cells in adulthood (Goodyear et al., 1999). In contrast, the fish ear and lateral line systems, which show robust and continuous regeneration of hair cells, relies on SuSCs or mantle cells for slow replacement of hair cells and on the availability of uncommitted multipotent stem-like cells for major reconstruction of the neuromast architecture (**Figure 6**). Importantly, all of our experiments were carried out using a 1h or 2h exposure to 10 μM CuSO_4_, which causes death of hair cells and other cell types, including UHCP and SuSCs. Thus, many cell types have to be replaced and we are likely examining only one of the potential pathways for replacing hair cells in neuromasts.

**Figure 6.**
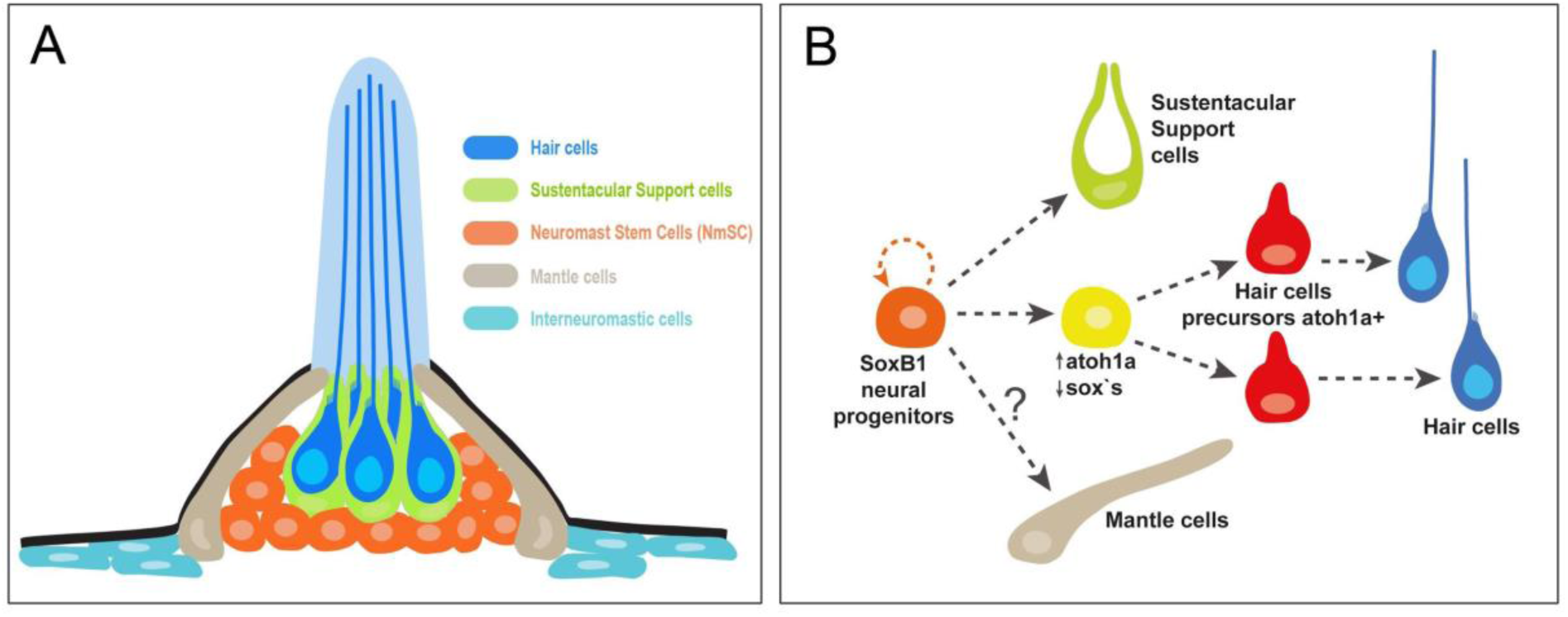
Proposed mechanism underlying different cell type regeneration in the pLL neuromast. (A) Cartoon representing the architecture and morphology of the different neuromast cell types described in this work. (B) Proposed lineages underlying each cell type regeneration after different damage level. The Neuromast Stem Cells (NmSC, orange) that express SoxB1 family proteins, self-renew and generate new Sustentacular Support Cells (SuSC, green), Unipotent Hair Cell mitotic Precursors (UHCP, yellow) and probably also Mantle cells (grey; not clarified in this study). Mitotic precursors that express *atoh1a* differentiate to mature hair cells (blue).

In the present article we propose an architectural map of the cell types of the neuromast. In order to compare our observations with published articles with a similar aim, we generated a comparative table showing the different cell types, their position within the neuromast, and the markers they express (**Table 1**). Following the new data obtained by us and other groups, it seems necessary to reach a consensus of the cell type diversity present in the neuromasts of the lateral line.

**Table 1.** Description of the different cell type populations in the neuromast. Comparison between our proposed cell type populations (left column) within the neuromast and published work by other groups (top arrow).

|  | Viader-Illargués et al. 2018 | Lush et al. 2019 | Romero-carvajal et al. 2015 | Thomas & Raible, 2019 | Seleit et al. 2018 (Medaka) | This work |
| --- | --- | --- | --- | --- | --- | --- |
| **Hair Cells** | SqEt4:GFP, Myo6b: actin-GFP | clusters 1,2 | SqEt4:GFP, *atoh1a*:dTom | brn3c:GFP, myo6b:GF<sup>FP</sup>, FM 1-43FX | Tg(Eya1:EGFP) | brn3c:GFP, Acetylated tubuline |
| **UHCP (Unipotent Hair Cell Progenitor, transient cells)** | not described | clusters 2,4: Proliferating Support Cells | *atoh1a*:dTom: AP Cells that regenerate | sost:nlsEos: some DV cells that regenerate HC | not described | *atoh1a*:dTom regenerative context |
| **SuSC (Sustentacular Support Cells)** | SqGw57a, Tg(Sox2: GFP): non-sensory sustentacular supporting cells | clusters 7,8,9,14: central supporting cells | not described | not described | Tg(Eya1:EGFP) | most Tg(HEs8:GFP) central and drop shape cells |
| **NmSC (Neuromast Stem Cells)** | SqGw57a, Tg(Sox2: GFP): non-sensory sustentacular supporting cells | clusters 3,10,11,12,13: DV pole and AP pole support Cells | DV Cells, Some AP cells | tnfsf10l3:nlsEos, sost:nlsEos: AP Cells and some DV Cells | neuromK8:H2B-EGFP, Tg(K15:H2B-RFP) | Sox2 & Sox3 cells, some Tg(HEs8:GFP) peripheral and round shape cells |
| **Mantle Cells** | Alpl:mCherry: Quiescent SC | clusters 5,6: Peripheral cells, Long-term SC | SqEt20:GFP: Long-term SC, Quiescent SC | sfrp1a:nlsEos: Long-term SC, Quiescent SC, AP Cells that not regenerate | neuromK8:H2B-EGFP: Long-term SC, Quiescent SC | some Sox2 & Sox3 cells |

### Sox2 and sox3 maintain a mitotically active population of neuromast stem cells

Using immunofluorescence, we observed the expression of Sox2 and Sox3 proteins in most cells of the migratory primordium, interneuromastic cells, pre- and pro-neuromasts, and neuromasts of the anterior and posterior lateral line systems, suggesting that both genes are required for proper early lateral line development, as is the case in the zebrafish otic placode (Gou et al, 2018a). In mature neuromasts, some Sox2_+_ and Sox3_+_ cells were also proliferating, indicating a low self-renewing process under steady state conditions. Our cell fate analysis of these proliferative cells after a short (2 h) and a long (24 h) BrdU pulse assay showed that all of the BrdU-labeled cells underwent a second division several days later, and almost half of these cells underwent another division one week later, showing slow self-renewal behavior, typical of stem cells. Also, under a damage-regeneration context, many Sox3_+_ progenitors retained this behavior of a slow cell cycle, indicative of self-renewal and the preservation of a stem cell niche. Recently, Lush et al. (2019) has shown by single-cell RNA-Seq that both Sox2 and Sox3 are expressed in almost the same population of neuromast cells. Therefore, we propose the definition of Neuromast Stem Cells (NmSCs) as these multipotent progenitors that reside within the base of the neuromast, morphologically different from sustentacular supporting and mantle cells, and that express Sox2 and Sox3 (**Figure 6**).

### Behavior of progenitor cells after repeated cycles of damage and regeneration

After damage by CuSO_4_ or neomycin, hair cell recovery in the zebrafish lateral line occurs after about 48 h (Harris et al., 2003; Hernández et al., 2007; Ma et al, 2008; Wibowo et al., 2011; Mackenzie and Raible, 2013; Cruz et al., 2015). Here, we demonstrate that this regenerative capacity is likely to be permanent and inexhaustible in the larvae as hair cells continue to be replaced after repeated injury. Other authors observed similar results in adult fish using neomycin treatment (Pinto-Texeira et al. 2015; Cruz et al. 2015). The maintenance of regenerative capacities in adults and larvae indicates that there is a nearly unlimited potential for restoration of lost hair cells that can only be explained by the presence of NmSCs in the zebrafish lateral line.

Single-cell RNA-Seq analysis of neuromasts has shown that both Sox2 and Sox3 are expressed in a few hair cells (Lush et al., 2019). We found that, in steady state, less than 7% of hair cells showed detectable levels of Sox2 protein (data not shown) while, during regeneration, this percentage increased to 50% and reached 100% when larvae were subjected to repeated rounds of injury. The latter finding likely reflects the overlap of consecutive cell fate transitions during regeneration, given that all hair cells present after repeated cycles of damage and regeneration are generated rapidly and directly from neuromast progenitors that continuously express Sox2. In fact, when we analyzed the expression of Sox2 within neuromast cells that incorporated BrdU after injury, almost all cells that proliferated also expressed this marker. In addition, we found that Sox2_+_ cells that divided during the BrdU pulse increased in proportion with consecutive rounds of damage-regeneration. This finding is similar to what was observed by Jaques et al. (2014), where stimulating proliferation within the lateral line through the activation of the Wnt/β-catenin pathway, increased the number of Sox2_+_ cells within the neuromast.

Our results provide further evidence that explains the alternative modes of regeneration that hair cells can undergo in neuromasts, as has been observed by other authors (Ma et al., 2008; Cruz et al., 2015; Romero-Carvajal et al., 2015; Lush et al. 2019; Viader-Llargués et al. 2018; Thomas and Raible, 2019). Indeed, after mild damage, the regenerated hair cells mostly derive from committed UHCP cells (video S3), located centrally in the neuromast that divide once to produce two hair cells and maintain the hair cell polarity (UHCP/*atoh1a*+ cells; Wibowo et al., 2011; or the *fgfr1a*+ cells; Lee et al., 2016; or the antero-posterior cells in Romero-Carvajal et al., 2015; or the clusters 2 and 4 in Lush et al., 2019, summarized in Table 1). With more severe injury or after repetitive rounds of injury, these unipotent hair cell progenitors are depleted and the organ must resort to the NmSCs, which can proliferate and regenerate all the components of a functional neuromast (**Figure 6B**).

### Functional analysis of *sox2/sox3* genes

It has been proposed that, in the zebrafish inner ear, *sox2* is necessary for hair cell survival and for the transdifferentiation of support cells into hair cells during regeneration (Millimaki et al., 2010). During development, loss-of-function analysis using *sox2* and *sox3* single mutants and morpholinos showed no overt abnormalities in embryo morphology, similar to what was found previously (Kamachi et al., 2008; Okuda et al., 2010). However, simultaneous knock-out and knock-down of both genes showed major defects in the lateral line development, with problems in protoneuromast formation, pro-neuromast deposition and hair cell number. Our finding could reflect that, in zebrafish, *sox2* and *sox3* have redundant roles in lateral line development as is the case for the establishment of normal otic and epibranchial tissues (Gou et al., 2018a). Interestingly, the functions of both genes are distinct in the inner ear, where *sox2* has a role in sensory development whereas *sox3* has a neurogenic role (Gou et al., 2018b).

It has been shown that *sox21* counteracts the activity of *soxb1* genes in chicken (Sandberg et al., 2005) and human glioma cells (Ferletta et al., 2011). In the zebrafish lateral line, knock-down of the *sox21a* gene results in increased cell death and failure in protoneuromast formation within the pLL primordium, a decreased in primordium migration speed and a reduction in the number of hair cells per neuromast, as well as in some cases abnormal deposition of the neuromasts (Ariza-Cosano et al., 2014). Because these phenotypes partially phenocopy the effect of blocking FGF signaling, Ariza-Cosano et al. (2014) concluded that *sox21a* might regulate the FGF pathway in the migratory primordium. In our experiments using double morphants, we did not observe a reduction in the number of primordium cells, possibly due to the accumulation of cells that were not deposited. We also detected a slight increase in cell death and impaired protoneuromast formation within the migrating primordium, delayed neuromast deposition and less hair cells within neuromasts. Altogether, our data suggest that *sox2* and *sox3* genes could play more specific roles in neuromast cell determination, while FGF signaling could play a role in primordium migration, and the frequency of pro-neuromast deposition (Durdu et al., 2014). A strong interaction could exist between the migration and deposition process, with participation of FGF signaling, and the determination of the cell types in the neuromast, where *sox* gene function becomes more relevant.

The relationship between Sox2 and cell proliferation has been widely shown in cancer cells and adult neurogenesis, a role related with its expression into stem-like cells (Ferri et al 2004; Episkopou 2005; Stolzenburg 2012; Piva et al 2014). Within the neuromast, few cell divisions (all of them expressing *sox2* and *sox3*) are found in a quiescent stage. However, after hair cell ablation by copper injury, cell divisions increase substantially and more than 70% of these proliferating cells also express *sox2* and *sox3* genes, some of them probably NmSCs. Functional analysis using the electroporation of Morpholinos against these genes specifically into injured neuromasts blocked cell division and confirmed the essential and active role that *sox2* and also *sox3* have in the triggering of cell proliferation in a regeneration context.

In conclusion, the results described here reveal that *sox2* and *sox3* are essential for the development of the zebrafish lateral line. We also generate a new classification that more accurately describes the different cell populations that reside in the lateral line neuromast, including the proposed SuSCs, with a strong ligation with the surrounding hair cells, and the NmSCs, as well as a description of the events that take place during its two types of regeneration (**Figure 6**). These findings further demonstrate the potential of the zebrafish lateral line as a powerful model for the understanding of hair cell development and regeneration.

## ACKNOWLEDGMENTS

We thank Drs. Pavla Navratilova and Thomas Becker for *Tg(Bglobin/HEs7:GFP)* strain, Dr. Steve Wilson for Sox3 antibody and to Drs. Joaquim Wittbrodt and Virginie Lecaudey for the time-lapse set-up. We are also thankful to Drs. Alain Ghysen and Christine Dambly-Chaudière for the set-up for electroporation and valuable comments and Drs. Lázaro Centanin and Leonardo Valdivia for critical reading of the manuscript. We thank the Aquatics Facility at the CRG for the fish care and Florencio Espinoza and Juan Silva for technical help.

---

**Figure Supplementary 1.**
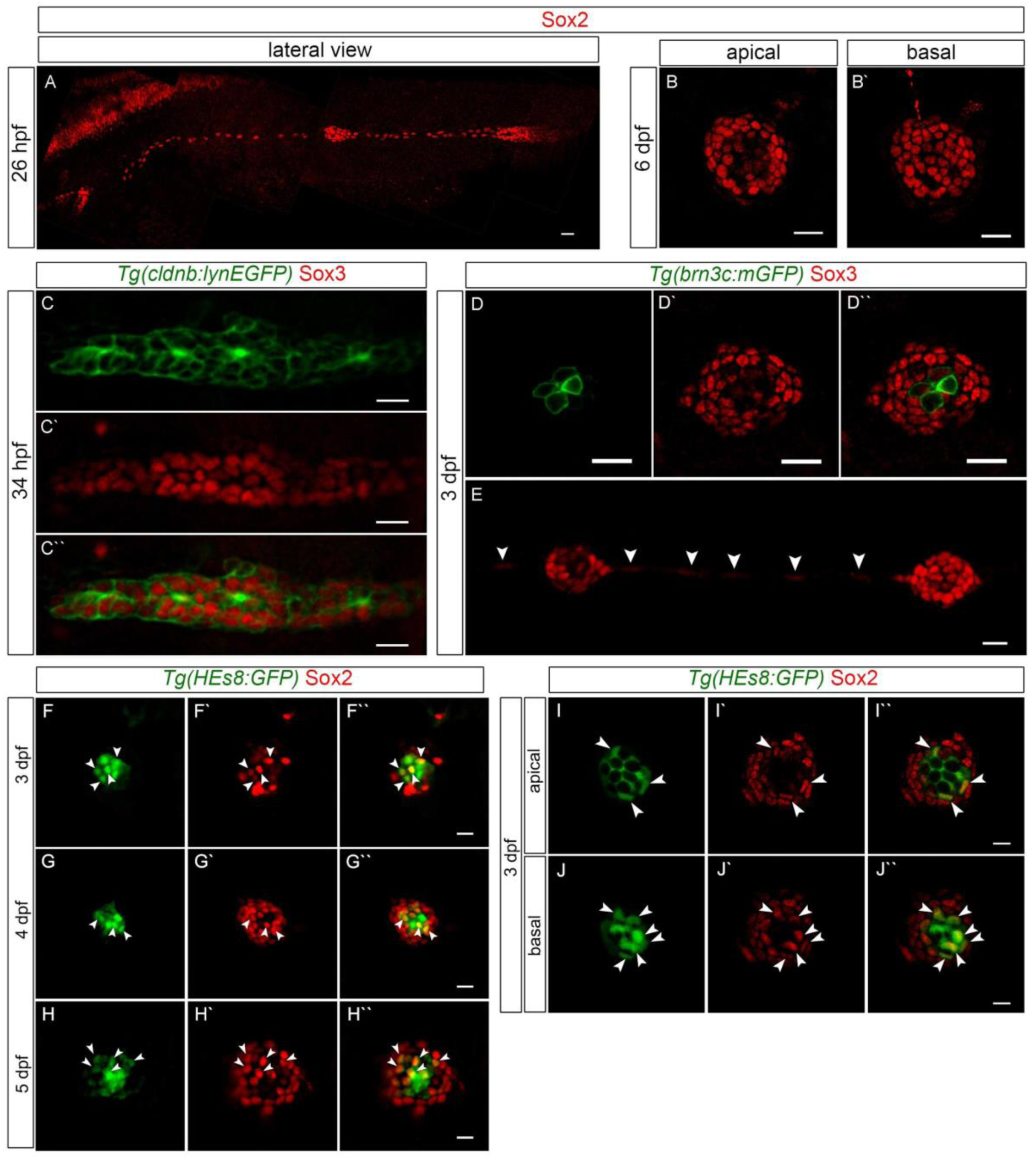
Expression of Sox2 and Sox3 in the primordium, neuromasts and interneuromastic cells of the posterior lateral line. (A) Immunofluorescence of Sox2 in the migratory primordium and proneuromast of pLL at 26 hpf and (B, B’) in neuromasts of the pLL at 6 dpf. (C-C’’) Confocal images of a migratory primordium at 34 hpf in a *Tg(cldnB:GFP)* transgenic embryo showing the presence of Sox3 in most of the cells. (D-D’’) Confocal images of neuromasts at 3 dpf in *Tg(brn3c:mGFP)* transgenic larvae, expressing Sox3 (red) in a population of cells that are not hair cells (green). (E) Two neuromasts of 3 dpf wt larvae expressing Sox3 in their cells and in the interneuromastic cells (arrowsheads). (F-H’’) Confocal immunofluorescence of Sox2 protein in 3 dpf, 4 dpf and 5 dpf *Tg(HEs7:EGFP)*. (I,J) Confocal immunofluorescence of Sox3 protein in 3 dpf *Tg(HEs7:EGFP)* larvae, with an apical view (I) and a basal view (J) of a neuromast. Scale bars = 10 μm.

**Figure Supplementary 2.**
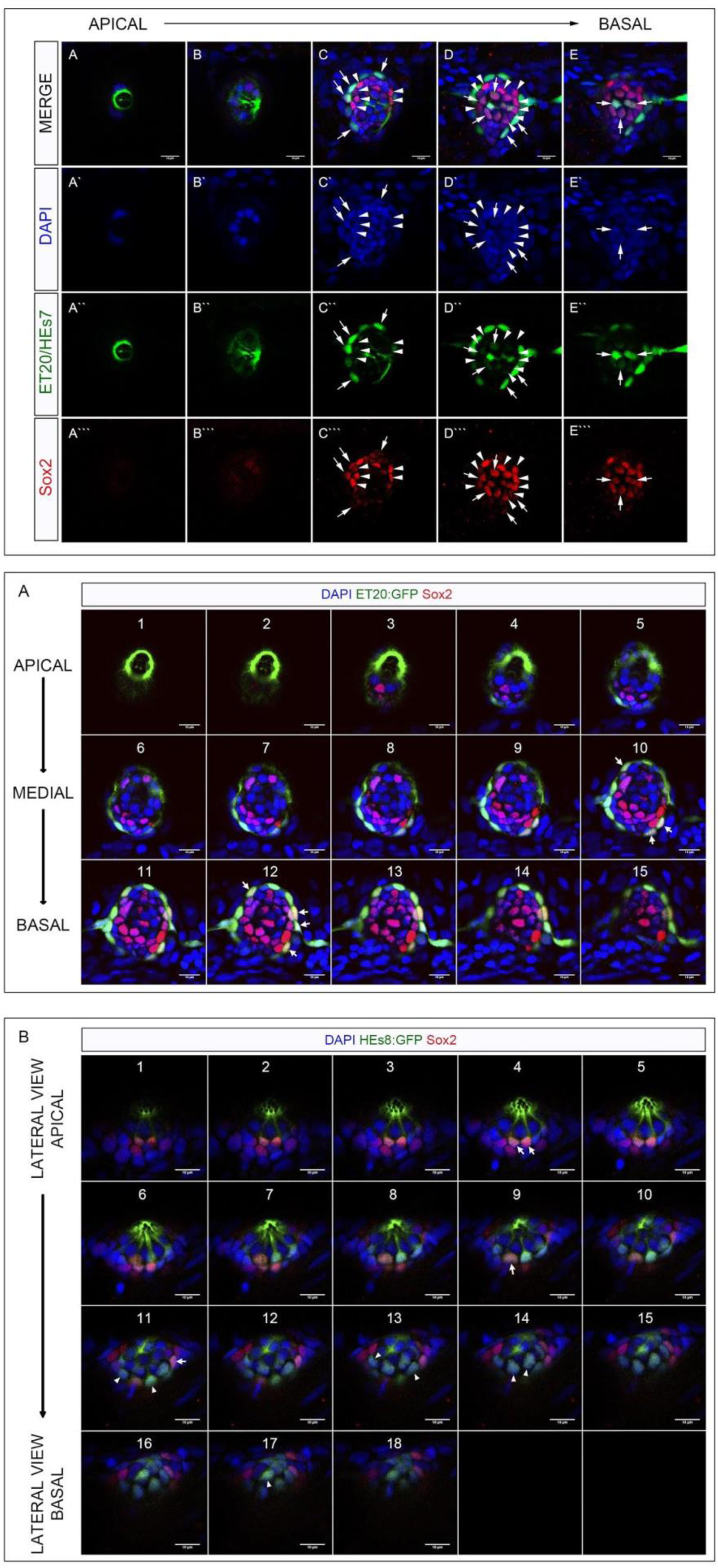
Distribution of mantle cells, support cells and Sox2_+_ cells in the neuromasts. (A-E’’’) Serial stacks of confocal images from apical to basal neuromast of 4 dpf double transgenic larvae *Tg(ET20:EGFP;HEs7:EGFP)* with immunofluorescence of Sox2 (red) and DAPI nuclear marker (blue). These images show that the GFP_+_ accessory cells (mantle and support cells) express the stem cell marker Sox2 (arrows) and also the expression of Sox2_+_ cells without colocalization with GFP_+_ cells (arrowheads). (F) Serial stacks of confocal images from apical to basal neuromast of 4 dpf in a transgenic larvae *Tg(ET20:EGFP)* with immunofluorescence of Sox2 (red) and DAPI nuclear marker (blue). The GFP_+_ mantle cells colocalize with the stem cell marker Sox2 (arrows) at the periphery of neuromasts with the characteristic half-moon shape. Most of the Sox2+ cells that do not colocalize with GFP_+_ cells are localized more central in the neuromasts and had a spherical shape. (G) Serial stacks of confocal images from a lateral-apical view to lateral-basal view neuromasts from 4 dpf transgenic larvae *Tg(HEs7:EGFP)* with immunofluorescence of Sox2 (red) and DAPI nuclear marker (blue). The GFP_+_ support cells that colocalize with Sox2_+_ cells are indicated with arrows and had the characteristic shape engulfing the hair cells (indicated by their large nuclei). The few support cells that do not colocalize with Sox2 are indicated with arrowheads. Scale bars = 10 μm.

**Figure Supplementary 3.**
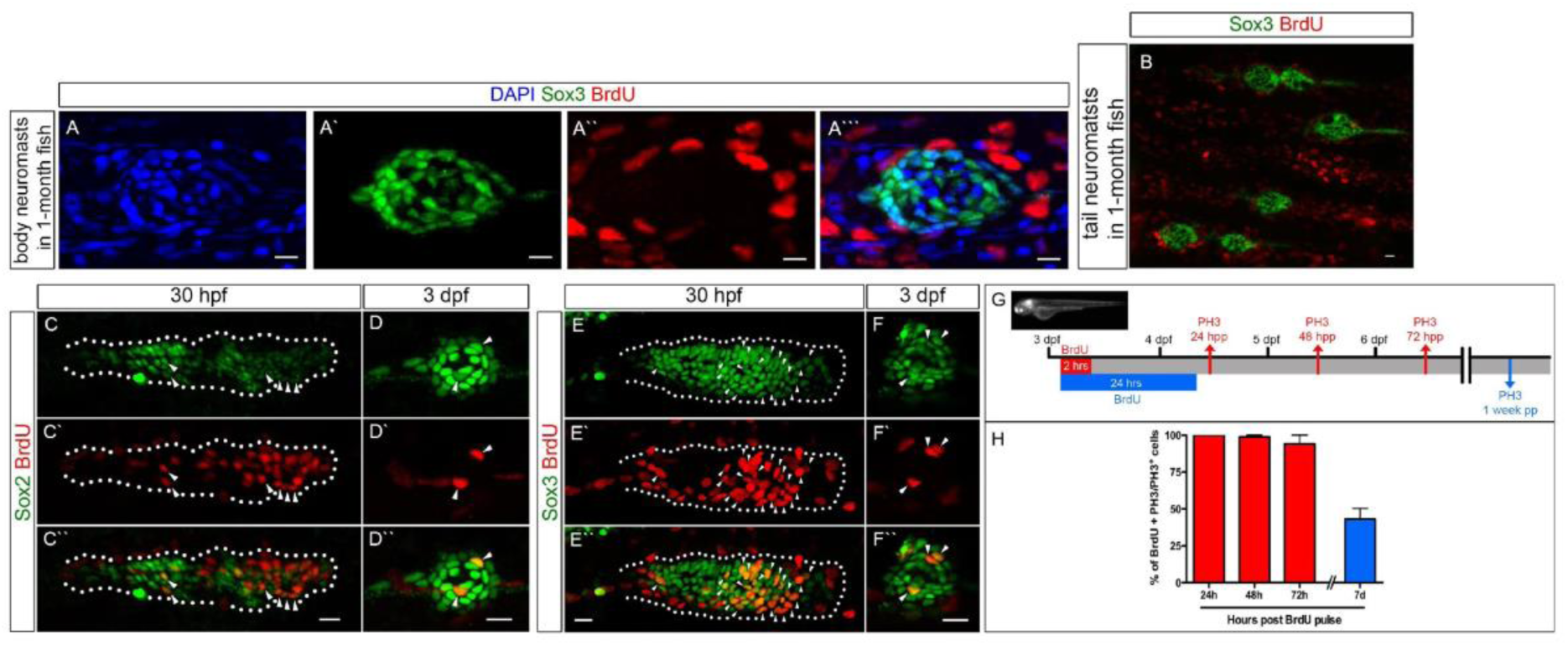
Proliferation of Sox2 and Sox3 positive cells in the primordium and neuromast and self-renewal cells in the posterior lateral line. (A) Confocal images of a body neuromast in 1-month-old fish. The Sox3+ cells are labeled by immunofluorescence; the proliferative cells after a 24 h pulse of BrdU are labeled in red and the DAPI staining marks the nuclei in blue. (B) Confocal images of BrdU+ proliferating cells (red) and Sox3+ cells (green) in the tail fin neuromasts of a 1-month-old fish after a 24 h pulse of BrdU. (C-F) Wt larvae at 28 dpf and 3 dpf were incubated for 2 h in BrdU and fixed for immunofluorescence detection. Confocal images of double immunolabeling for Sox2 positive cells (green) and BrdU positive cells (red) in the migratory primordium of embryos at 30 hpf and in a neuromast of a larvae at 3 dpf (C-D). The cells that co-express both markers are indicated with arrowheads. (E-F) Confocal images of double immunolabeling for Sox3-positive cells (green), and BrdU positive cells (red) in the migratory primordium (30 hpf embryo) and neuromasts (3 dpf larvae). The cells that co-express both markers are indicated with arrowheads. (G) Scheme showing the two BrdU pulse-fix time course used and the fix times for PH3 immunofluorescence. 3 dpf larvae were incubated 2 h in BrdU and fixed 24, 48 and 72 h later (red) and 3 dpf larvae were incubated for 24 h in BrdU and fixed 1 week later (blue). (H) Almost all cells labeled with a short pulse have divided (PH3 label, red bars) and one-week post long pulse, 40% of the cells have divided (PH3 label, blue bar). Scale bars = 10 μm.

**Figure Supplementary 4.**
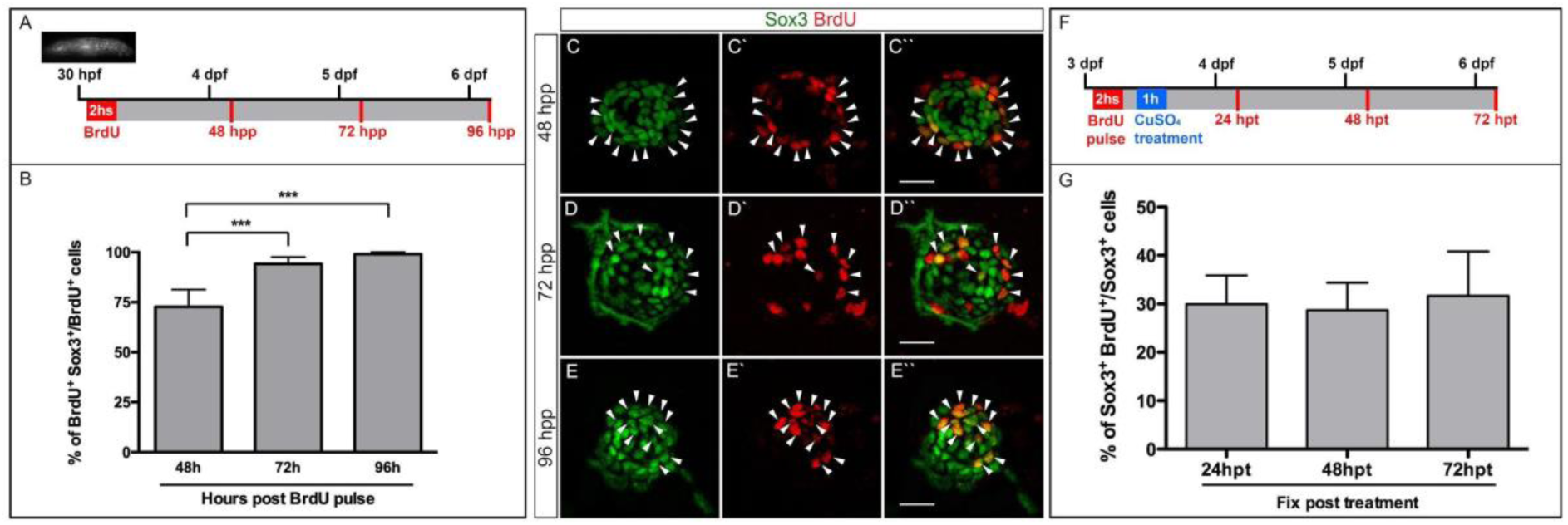
Permanence and location of Sox3 proliferating cells in the primordium and neuromasts. (A) BrdU pulse-labeling scheme representing the experiment in which proliferating cells in the primordium of 30 hpf larvae were labeled with BrdU pulses and the colocalization with Sox3 positive cells in mature neuromasts was observed at different chase times (arrow head, C-E). (B) The quantification shows that most of proliferative cells in the primordium express the Sox3 neural progenitor marker in the mature neuromasts and this percentage is increasing significant post BrdU pulse (P<0.01). (F) Scheme representing the BrdU pulse before the incubation with copper sulfate and the different chase times. (G) The graph shows that almost 30% of all the Sox3_+_ cells retain the BrdU label at 24, 48 and 72 h post BrdU incorporation. Scale bars = 10 μm.

**Figure Supplementary 5.**
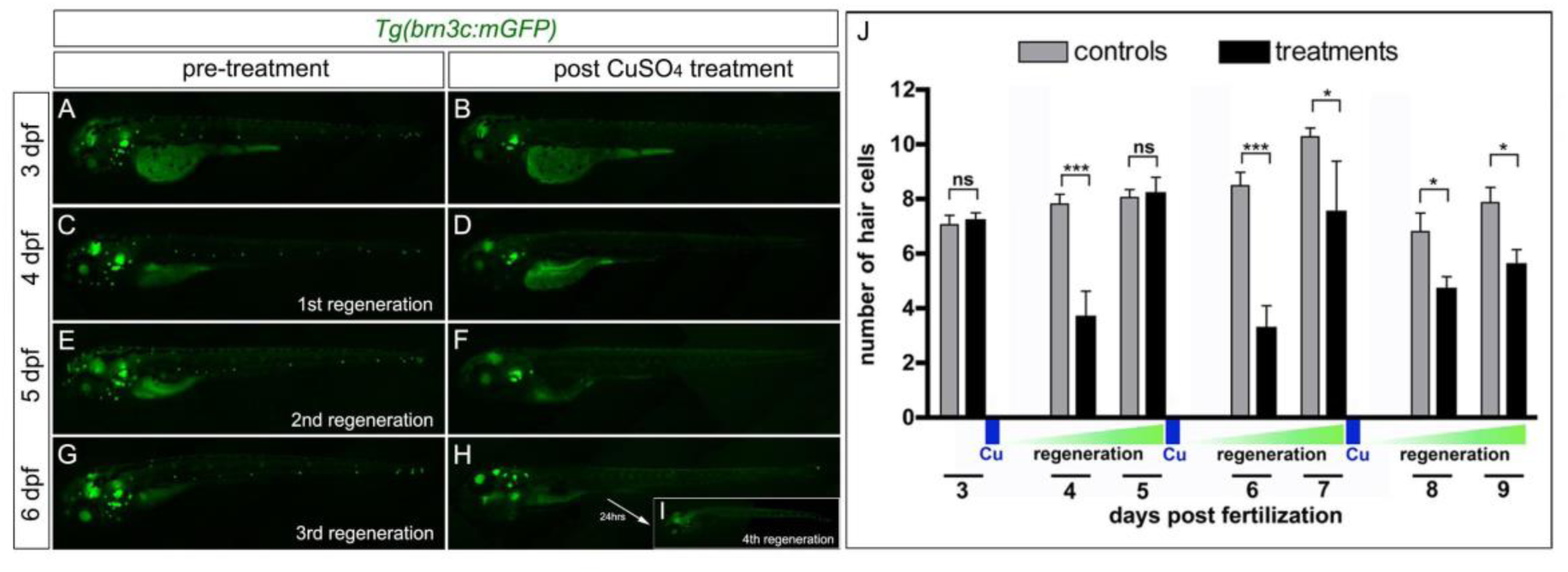
Hair cell regeneration cycles in the pLL of transgenic zebrafish larvae. Mature and regenerated hair cells are labeled by the *Tg(brn3c:mGFP)* transgene. (A) A 3 dpf larvae prior to the first treatment showing the discrete pattern of GFP-positive hair cells in the pLL. In (B, D, F, H) a complete elimination of hair cells after damage with copper sulfate is shown, assessed by the absence of GFP. Regenerated hair cells in the pLL 24 h after each treatment are shown in (C, E, G, I). Four consecutive cycles of regeneration were observed. (J) Labeled hair cells were counted daily and after each copper treatment in the 3 first neuromasts of the pLL. Partial recovery in the number was observed at 24 h after the first treatment (4 dpf), and a complete recovery of the hair cell population was evident after 48 h (5 dpf). At the second and third cycles of regeneration even if we observed a robust regenerative capacity (6 – 9 dpf), the number did not reach those for hair cells in control larvae; this holds true even 48 h after both treatments (7 and 9 dpf).

**Figure Supplementary 6.**
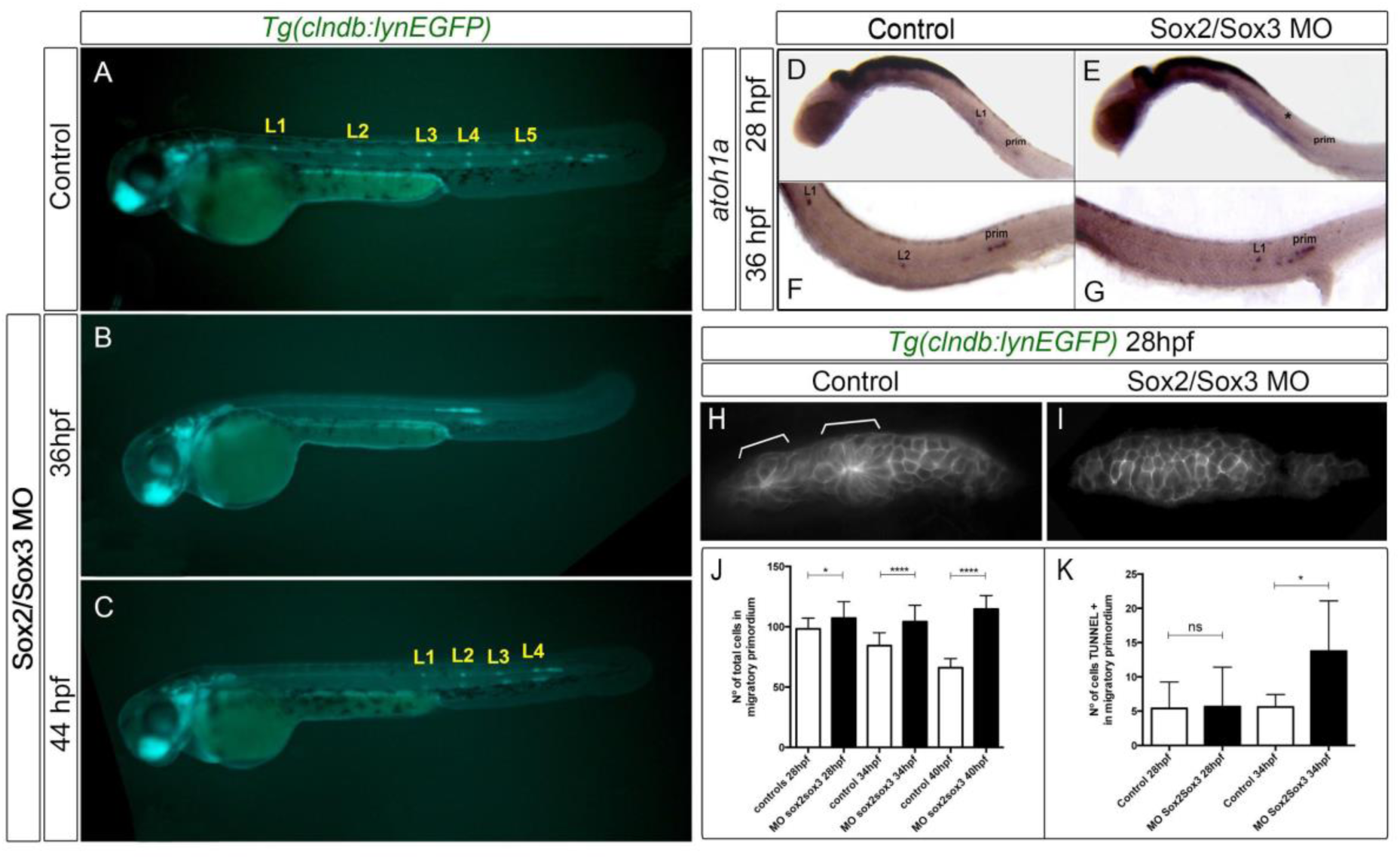
Normal deposit of pLL neuromasts requires the concerted action of SoxB1 family genes. (A-C) 46 hpf transgenic larvae *Tg(cldnB:lynGFP)* showing neuromast distribution (from L1 to L5). (A) Control larvae showing a normal pattern of deposited neuromast. (B, C) Sox2/Sox3 double morphants fail to deposit the neuromast at early stages. The migratory primordium finishes at the end of the tail, and deposited the neuromast in the last third of the tail observed at 44 hpf (C). (D-G) The RNA expression of *atoh1a* is absent in the 28 hpf double morphant larvae (E), compared with the wt larvae (D) and just observed at 36 hpf, when the migratory primodium start to deposit, delayed, the first proneuromast across the tail (G). At the same stage in control larvae the expression of *atoh1a* is observed in two of the proneuromasts (F). (H-I) At 28 hpf *tg(cldnB:GFP)* primordium shows two protoneuromasts at the trailing domain pre-forming the future neuromast (H, white brackets). sox2/sox3 double morphants fail to organize cells into protoneuromasts in the primordium, showing a general cellular disorganization (I). These morphology differences were observed in the number of total cells of the migratory primordium at different stages (J) and in the number of TUNEL positive cells at 28 hpf and 34 hpf (K).

**Figure Supplementary 7.**
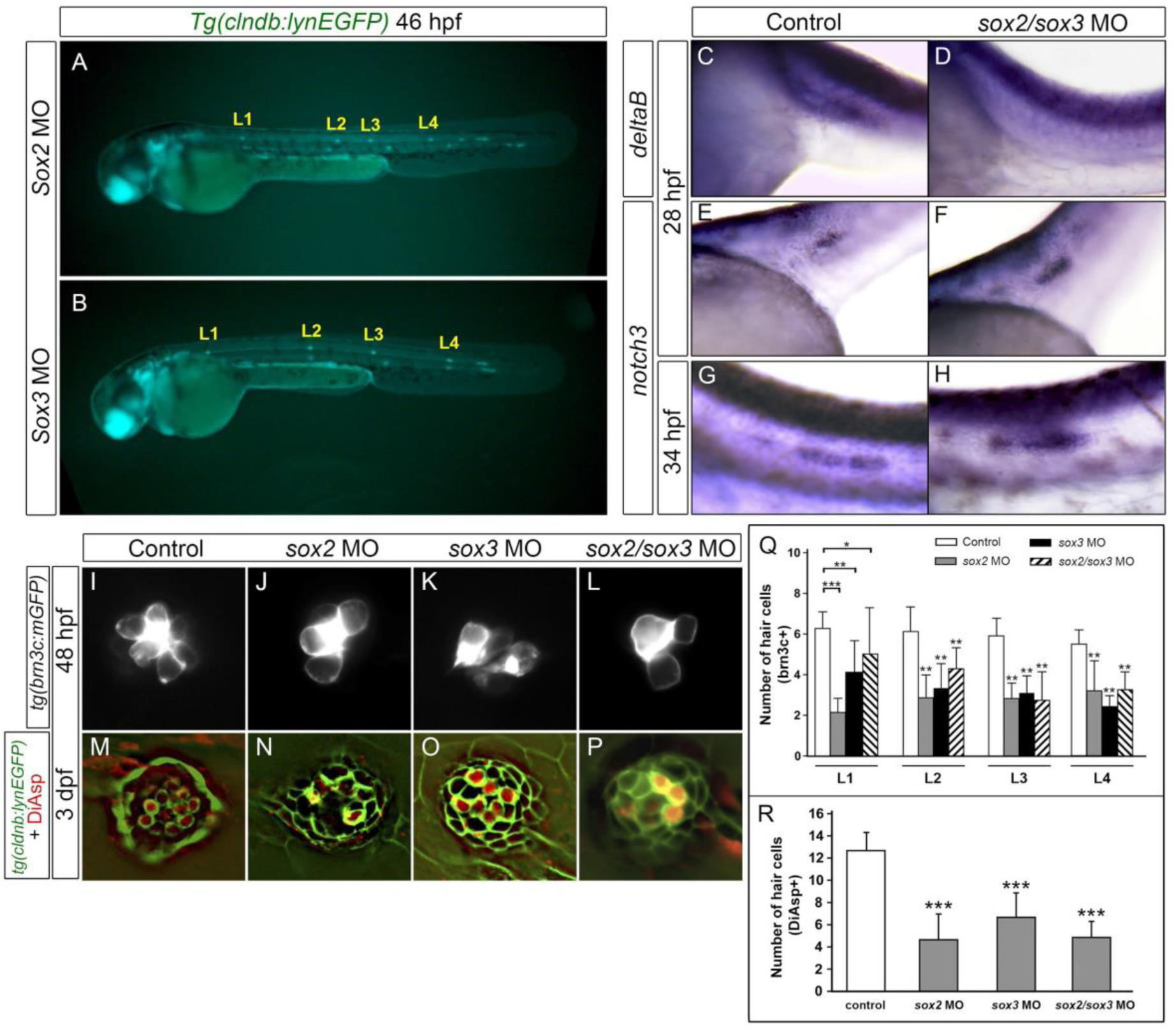
*sox2* and *sox3* knockdown result in changes in the expression patterns of *deltaB*, *notch3* and impairs pLL lateral line formation. (A-B) Normal deposition of proneuromasts in both *sox2* and *sox3* knockdown at 46 hpf in *tg(cldnB:GFP)*. (C–H) Lateral view of whole-mount in situ hybridizations for *deltaB* (C-D) and *notch3* (E-H) in control and double morphant embryos at 28 hpf and 34 hpf (anterior is to the left and dorsal is up). *deltaB* expression is lost in the morphant primordium at 28 hpf, while *notch3* shows a similar or slightly higher level when compared to control at 28 and 34 hpf. (I–P) 48 hpf transgenic embryos *tg(brn3c:GFP)* were used to analyze the number of hair cells in the pLL neuromast in single and double morphants for *sox2* and *sox3*. (Q) Quantitative analyses in the first four neuromasts (from L1 to L4) show differences in hair cell number between controls and the 3 different morphant conditions. These differences in hair cell numbers (assessed by DiAsp labeling) were maintained at 3 dpf. (R) Average number of hair cells in the first three neuromasts (from L1 to L3) at 3 dpf, showing the significant differences between each morphant and the control.

**Figure Supplementary 8.**
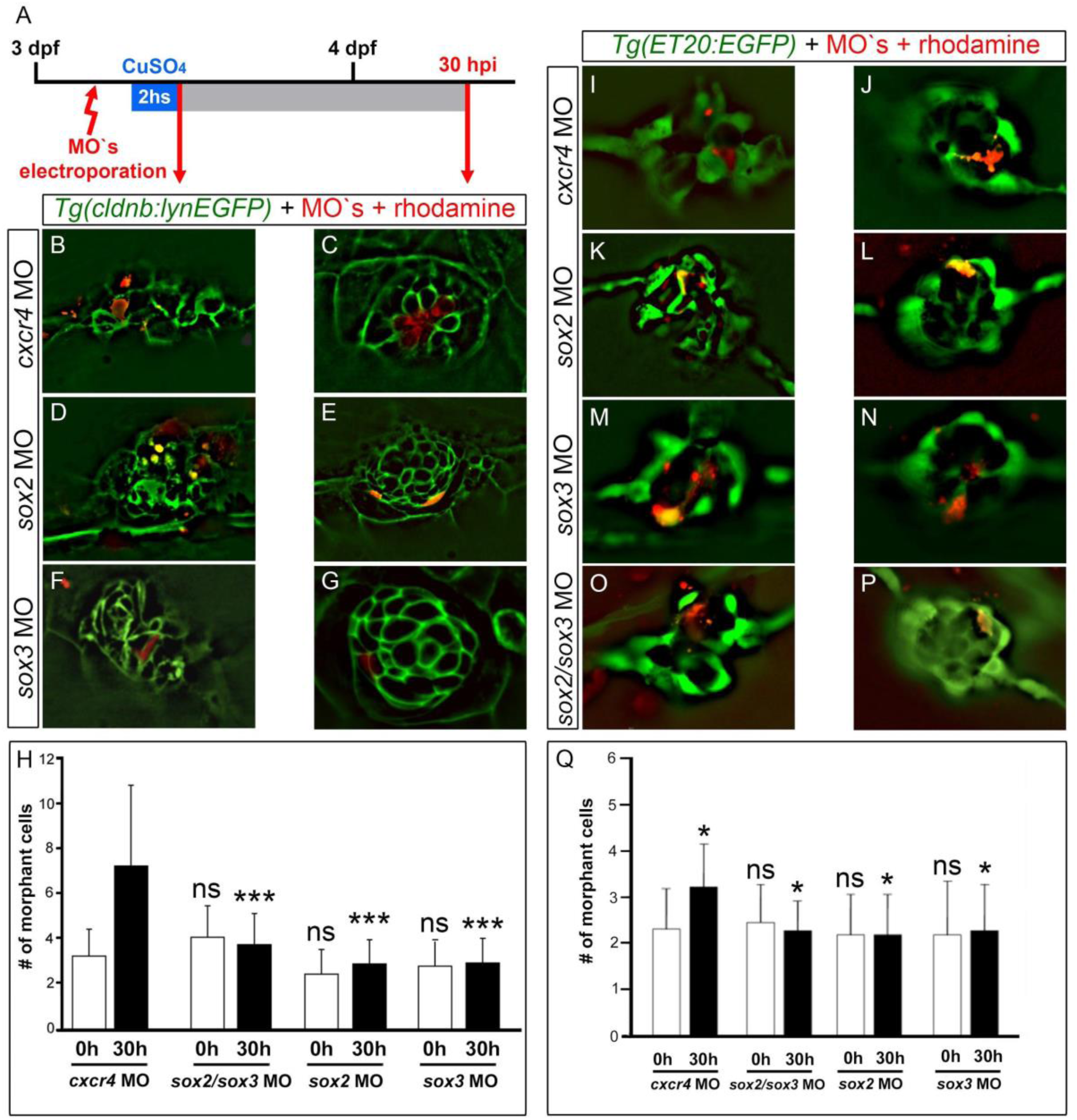
*sox2* and *sox3* are necessary for the proliferation and regeneration in pLL neuromasts. (A) Scheme representing the electroporation and regeneration experiments, where the neuromasts of the 3 dpf transgenic larvae *Tg(cldnB:GFP)* and *Tg(ET20:GFP)* were electroporated with rhodamine-tagged *cxcr4* MO (controls) and rhodamine-tagged morpholinos for *sox2*, *sox3* and *sox2/sox3*, after were incubated with CuSO_4_ for 2 h to eliminate the hair cells and fixed at 0 and 30 h post copper incubation (hpi). (B-G) *Tg(cldnB:GFP)* larvae, which express GFP in all neuromast cells were electroporated with different morpholinos and analyzed immediately before and after copper treatment (B, D, F 0 hpi, and C, E, G 30 hpi). The same analyses were performed for the *Tg(ET20:GFP)* larvae, which express GFP in the mantle cells of the neuromasts. (H,Q) Number of “morphant cells” (cells that incorporated the rhodamine-tagged morpholino) counted immediately before (0 hpi) and after copper treatment (30 hpi) in both transgenic lines. The graphs shows statistically significant differences in the amount of “morphant cells” (p<0.001 for *cldnB:GFP* and p<0.05 for *ET20:GFP*), indicating a proliferative inhibition on morphant neuromasts, thus blocking cell divisions, an essential step for NmSC to divide and regenerate new hair cells.

**Video S1.** Lateral reslice of confocal serial stack images of a double transgenic line *Tg(HEs7:EGFP;atoh1a:dTomato)* + DAPI to label the nucleus. The GFP label the Sustentacular Support Cells (SuSC) that completely surround the committed hair cells labeled by *dTomato*.

**Video S2.** 3D reconstruction of confocal serial stacks image of a double transgenic line *Tg(HEs7:EGFP;atoh1a:dTomato)* + DAPI to label the nucleus.

**Video S3. Hair cells differentiate from a mitotic precursor during regeneration.** Hair cell elimination after copper treatment and new hair cell formation from pre-mitotic *atoh1a*-expressing hair cell-precursors. Time-lapse confocal microscopy imaging of a double *Tg(brn3c:mGFP/atoh1a:dTomato)* transgenic neuromast immediately after copper treatment showing the complete depletion of hair cells and the posterior emergence of new hair cells from an *atoh1a+* hair cell-precursor who divides.

